# An apical protein, Pcr2, is required for persistent movement by the human parasite *Toxoplasma gondii*

**DOI:** 10.1101/2022.05.20.492694

**Authors:** Jonathan Munera Lopez, Isadonna F. Tengganu, Jun Liu, John Murray, Luisa F. Arias Padilla, Ying Zhang, Peter T. Brown, Laurence Florens, Ke Hu

**Affiliations:** Biodesign Center for Mechanisms of Evolution/School of Life Sciences, Arizona State University, USA; Department of Biology, Indiana University, USA; Berkgen Biopharmaceuticals, Shenzhen, China; Stowers Institute for Medical Research, Kansas City, USA; Department of Physics and Center for Biological Physics, Arizona State University, USA

## Abstract

The phylum Apicomplexa includes thousands of species of unicellular parasites that cause a wide range of human and animal diseases such as malaria and toxoplasmosis. To infect, the parasite must first initiate active movement to disseminate through tissue and invade into a host cell, and then cease moving once inside. The parasite moves by gliding on a surface, propelled by an internal cortical actomyosin-based motility apparatus. One of the most effective invaders in Apicomplexa is *Toxoplasma gondii*, which can infect any nucleated cell and any warm-blooded animal. During invasion, the parasite first makes contact with the host cell “head-on” with the apical complex, which features an elaborate cytoskeletal apparatus and associated structures. Here we report the identification and characterization of a new component of the apical complex, <u>P</u>re<u>c</u>onoidal <u>r</u>egion protein <u>2</u> (Pcr2). Pcr2 knockout parasites replicate normally, but they are severely diminished in their capacity for host tissue destruction due to significantly impaired invasion and egress, two vital steps in the lytic cycle. When stimulated for calcium-induced egress, Pcr2 knockout parasites become active, and secrete effectors to lyse the host cell. Calcium-induced secretion of the major adhesin, MIC2, also appears to be normal. However, the movement of the Pcr2 knockout parasite is spasmodic, which drastically compromises egress. In addition to faulty motility, the ability of the Pcr2 knockout parasite to assemble the moving junction is impaired. Both defects likely contribute to the poor efficiency of invasion. Interestingly, actomyosin activity, as indicated by the motion of mEmerald tagged actin chromobody, appears to be largely unperturbed by the loss of Pcr2, raising the possibility that Pcr2 may act downstream of or in parallel with the actomyosin machinery.

## INTRODUCTION

Cell movement allows cells to explore their environment, initiate physical interaction with other cells, and respond accordingly. Movement forms the basis of numerous processes such as embryonic development and the inflammatory response in animals. For the thousands of species of apicomplexan parasites that are responsible for many devastating diseases (including malaria and acute toxoplasmosis), movement is integral to their parasitic lifestyle. For instance, upon injection into the human host by a mosquito, parasite motility enables the *Plasmodium* sporozoites to migrate through the skin tissue, and cross a blood vessel wall into the bloodstream to be carried to the liver, where the sporozoites move out of the blood vessel and invade into liver cells [1]. Similarly, *Toxoplasma*, an extremely successful parasite that permanently resides in ∼ 20% of the people on Earth and can infect any nucleated cell and any warm-blooded animal, relies on motility to shove its way into a host cell, as well as to rapidly escape from a resource-exhausted host cell, disseminate, and reinvade into a fresh host [2-6].

During invasion, the parasite first makes contact with the host cell “head-on” with its apical complex, which features an elaborate cytoskeletal apparatus and associated structures that contain both structural and signaling proteins important for invasion [7-13]. As the parasite enters the host cell, it assembles a ring-like structure (the moving junction) at the entry point from proteins secreted from the rhoptries and the micronemes, which is critical for the invasion process [14-22]. As it invades, the parasite also extensively modifies the host cell’s plasma membrane at the entry point, and uses that modified membrane to form the parasitophorous vacuole that eventually completely envelops the intracellular parasite [16, 23-35]. The integrity of the host plasma membrane and viability of the host cell are preserved during invasion, thus allowing for subsequent replication cycles during which the parasites exploit host resources for proliferation. Parasite exit (i.e. egress) on the other hand, is lethal for the host cell [3, 7, 36-38]. During *Toxoplasma* egress, the combination of pore formation by lytic proteins secreted from the parasite and mechanical disruption due to parasite moving through the host cell membrane results in destruction of the host cell, and allows parasite dissemination to initiate the next round of the lytic cycle. Successive cycles of parasite invasion, replication, and egress lead to extensive tissue damage such as seen in toxoplasmic encephalitis and congenital toxoplasmosis. Therefore, parasite motility not only is required to initiate an infection, but also directly contributes to the pathogenesis of the disease.

Infectious apicomplexan parasites move by gliding on a surface, propelled by an internal cortical actomyosin-based motility apparatus. Over the years, many labs have characterized key proteins involved in parasite motility [2, 6, 7, 12, 13, 39-46]. The functions of these proteins can be largely explained in the context of a working model [47], in which the internal motor activity powers parasite gliding via the coupling of the actomyosin machinery with transmembrane adhesin complexes (more in discussion). Here we report the discovery of a new apical complex protein, <u>P</u>re<u>c</u>onoidal <u>r</u>egion protein <u>2</u> (Pcr2), which is required for persistent movement. The loss of Pcr2 results in an unusual motility phenotype.

Pcr2 is a novel protein located in the apical complex of the parasite. Knockout of Pcr2 results in stuttering parasite movement, a phenotype that has not been observed before. Calcium stimulated basal accumulation of mEmeraldFP tagged actin chromobody (actin-Cb-mE) is not blocked in the Pcr2 knockout parasite. This suggests that Pcr2 does not regulate motility through actin polymerization or actomyosin activity, as both have been shown to be necessary for the basal accumulation of actin-Cb-mE to occur [13, 48]. The abnormal motility of the Pcr2 knockout parasite is associated with significantly reduced efficiency of parasite egress, invasion, and destruction of host cells, indicating the importance of Pcr2 for the lytic cycle.

## RESULTS

### Identification of new candidate preconoidal proteins using immunoprecipitation

During invasion, the parasite initiates contact with the host cell with its apical complex [49-53]. The cytoskeletal apical complex contributes to parasite invasion both structurally and as a signaling center [7-13]. It is a striking cytoskeletal machine, formed of a truncated cone-like basket (the conoid) and a series of associated structures, including the intraconoid microtubules, the preconoidal rings and the apical polar ring (Figure 1A). Disruption of the conoid structure is linked to severely impaired parasite invasion [8, 10]. In *Toxoplasma*, the 14 conoid fibers, novel tubulin polymers shaped like folded ribbons, are organized into a left-handed spiral capped at the apical end by the preconoidal rings. The preconoidal rings have an intricate periodic structure. The left panel of Figure 1B shows the EM image of a preconoidal ring viewed end-on, which has an outer diameter of ∼ 220 nm and an inner diameter of ∼ 160 nm. The substructures packed regularly in the ring somewhat resemble blunt spikes (arrowheads). Dividing the length of the segment where the spikes are visible by the number of spikes suggests an average periodicity of ∼ 16 nm. Even after the conoid fibers are separated from the rest of the parasite cytoskeleton following detergent extraction and protease digestion, they often are retained around or remain attached to a preconoidal ring (Figure 1B, *middle* and *right* panels), indicating that the ring and conoid fibers are structurally integrated. It is therefore conceivable that the preconoidal rings form the organizing center of the conoid.

**Figure 1.**
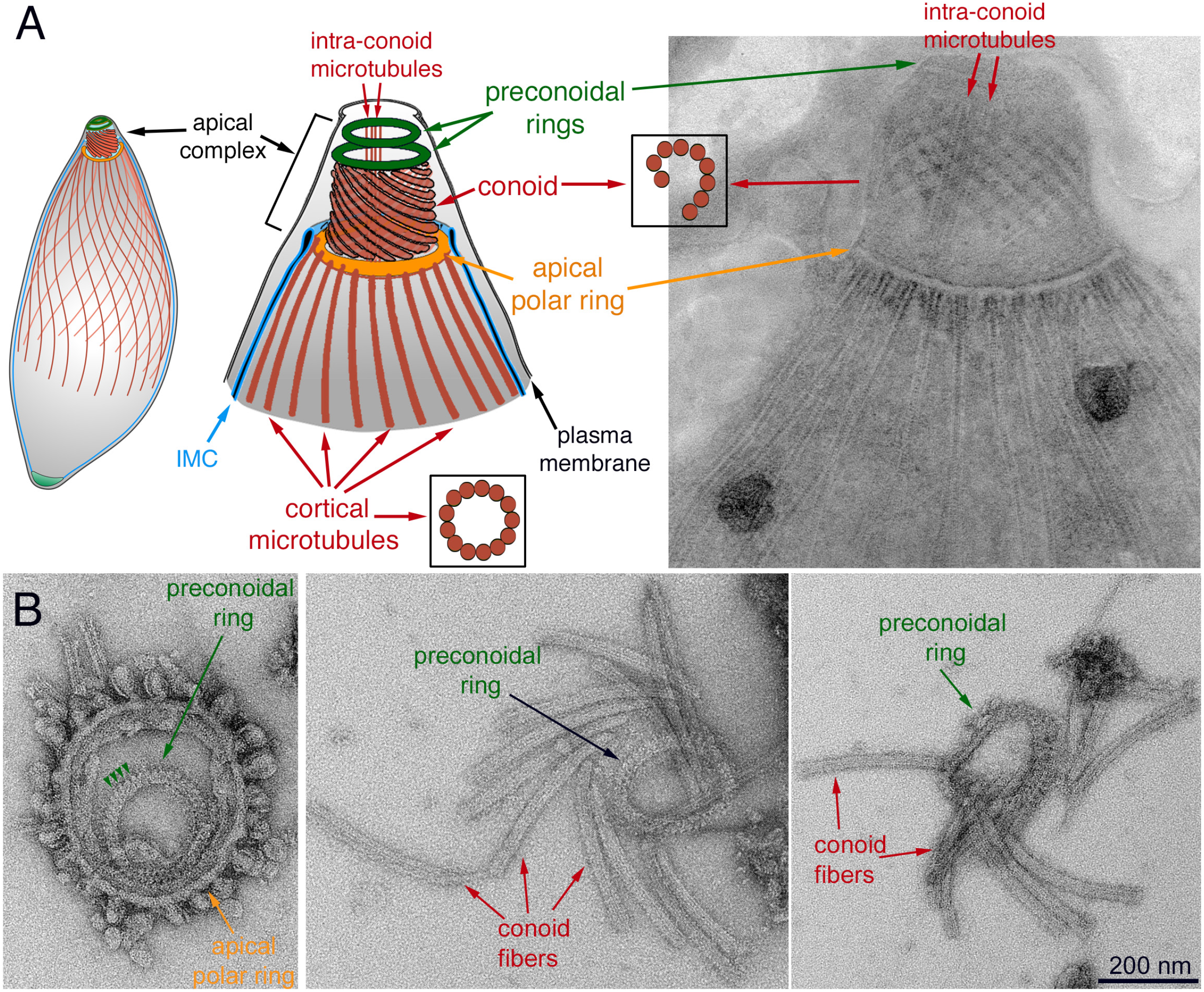
Structure of the apical complex in *Toxoplasma* **A**. *Left* and *middle:* Drawings depicting some of the tubulin-containing structures (red) in *T. gondii*, including the 22 cortical microtubules, a pair of intra-conoid microtubules, as well as the 14 fibers that make up the conoid. IMC: Inner Membrane Complex. *Right:* Transmission electron microscopy (TEM) image of a negatively stained TritonX-100 (TX-100) extracted parasite. **B**. TEM images of the apical parasite cytoskeleton negatively stained after detergent extraction and protease treatment. *Left:* end-on view of a parasite apical cytoskeleton. Most cortical microtubules and conoid fibers have detached, which allows a clear view of the preconoidal rings lying inside the apical polar ring. Arrowheads indicate the periodic “spikes” in the preconoidal ring. *Middle* and *right*: disassembled conoids with attached preconoidal rings.

Previously, we identified TgCentrin2 (CEN2), which localizes to the preconoidal rings and several other structures - the centrioles, peripheral annuli (also referred to as “apical annuli”), and the basal complex, and plays an important role in parasite invasion and replication [53-55]. To identify other potential preconoidal proteins, we performed immunoprecipitation (IP) analysis using GFP-Trap and lysate from a knock-in parasite line that expresses eGFP-CEN2 from the endogenous locus [55] (Figure 2A). <u>M</u>ultidimensional <u>P</u>rotein Identification <u>T</u>echnology (MudPIT) analysis identified with high confidence a number of proteins known to be localized to CEN2 containing structures (e.g., the centrioles, peripheral annuli, and the basal complex) (Table S1). In addition, three of the hypothetical proteins enriched in the IP fraction are localized to a structure that is apical and smaller in diameter compared to the conoid (Figure 2B-C), as predicted for proteins located at the preconoidal region. These proteins were thus named Pcr1 (TGGT1_274160), Pcr2 (TGGT1_257370), and Pcr3 (TGGT1_231840). The MS spectral counts of the three proteins per replicate are ∼ 99 (Pcr1), 228 (Pcr2), and 60 (Pcr3), respectively. The difference in these MS counts could be affected by the nature of the interaction between the bait and the pulled-down protein and the abundance of individual proteins, as well as by the length of the protein, among other factors. Pcr2, for instance, is the largest protein of the three, which could contribute to the higher spectral count. During our process of characterizing these proteins, a LOPIT (Localization of Organelle Proteins by Isotope Tagging) proteomics screen [56] confirmed the apical localization of Pcr1. We decided to focus on Pcr2, which was predicted to be strongly fitness conferring (phenotype score = -4.43) in a large-scale CRISPR-based screen [57], but had not been characterized.

**Figure 2.**
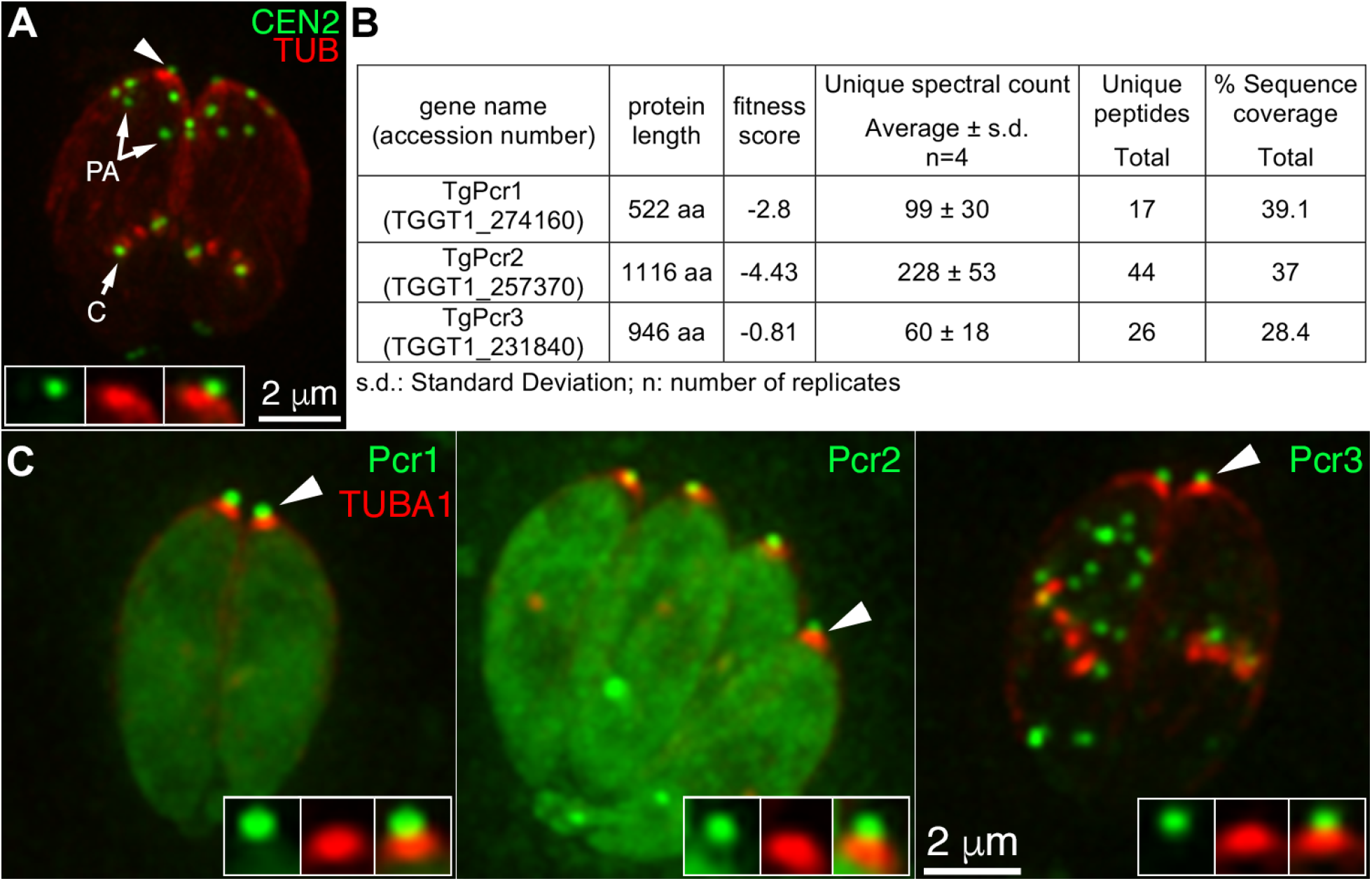
Identification of candidate apical complex proteins via immunoprecipitation using GFP-Trap and *eGFP-CEN2* knock-in parasites **A**. Deconvolved wide-field images of intracellular *eGFP-CEN2* knock-in parasites [55] expressing mCherryFP tagged tubulin (red). Insets (shown at 2X) include the preconoidal region (arrowhead). PA: peripheral annuli; C: centrioles. **B**. Table showing the protein length, fitness score, average unique spectral counts, peptide counts, and sequence coverage for TGGT1_274160, TGGT1_257370, and TGGT1_231840, identified by MuDPIT in 4 replicates of immunoprecipitation using GFP-Trap and *eGFP-CEN2* knock-in parasites. See Table S1 for the complete list of identified proteins. **C**. Deconvolved wide-field images of intracellular parasites expressing mCherryFP tagged tubulin (red) and mEmeraldFP (green) tagged Pcr1 (TGGT1_274160), Pcr2 (TGGT1_257370), or Pcr3 (TGGT1_231840) with expression driven by a *T. gondii* tubulin promoter. As predicted for preconoidal proteins, Pcr1, Pcr2, and Pcr3 are localized to a structure (green, insets) that is apical and smaller in diameter than the conoid (red, insets). Insets (shown at 2X) include the preconoidal region (arrowheads).

Pcr2 is a novel protein. Blast search against the database of Non-redundant protein sequences (nr-NCBI), yielded no hit with E value more significant than 0.05 outside the *Sarcocystidae* family, or conserved domains with E value more significant than 1e-5. Alpha-fold [58] predicted with confidence (pLDDT>70) four extended alpha-helical regions that likely form a coiled-coiled structure for protein-protein interaction, with no predicted structure for much of the rest of the protein (Figure S1).

To determine the endogenous localization of Pcr2, we generated an endogenous 3’ mNeongreenFP tagged line (*Pcr2-mNeonGreen* 3’tag) by single-crossover homologous recombination, and a knock-in line (*mEmeraldFP-Pcr2* knock-in) by double-crossover homologous recombination (see below). The FP-Pcr2 fluorescence is concentrated at the apical end of the mature parasite (Figure 3A). FP-Pcr2 fluorescence is retained after extraction with Triton-X100 (TX-100), and colocalizes with the preconoidal labeling of CEN2 (Figure 3B), indicating that, similar to CEN2 [53-55], Pcr2 is a stable component of the apical complex. The recruitment of Pcr2 to the preconoidal region is detectable when the nascent daughters first emerge in the mother, marked by the duplication of the spindle pole, the assembly of the tubulin-containing cytoskeleton that includes the conoid, and the growing cortical microtubules [52-54, 59](Figure 3C-D). In a small fraction of cells, a Pcr2-containing spot is also found close to the basal end of the parasite (Figure 3D, top row, arrowhead). The significance of this additional Pcr2 focus is unknown as it does not appear to consistently associate with any recognizable cellular structures. Pcr2 remains at the apex in invading parasites (Figure S2 and Video S1).

**Figure 3.**
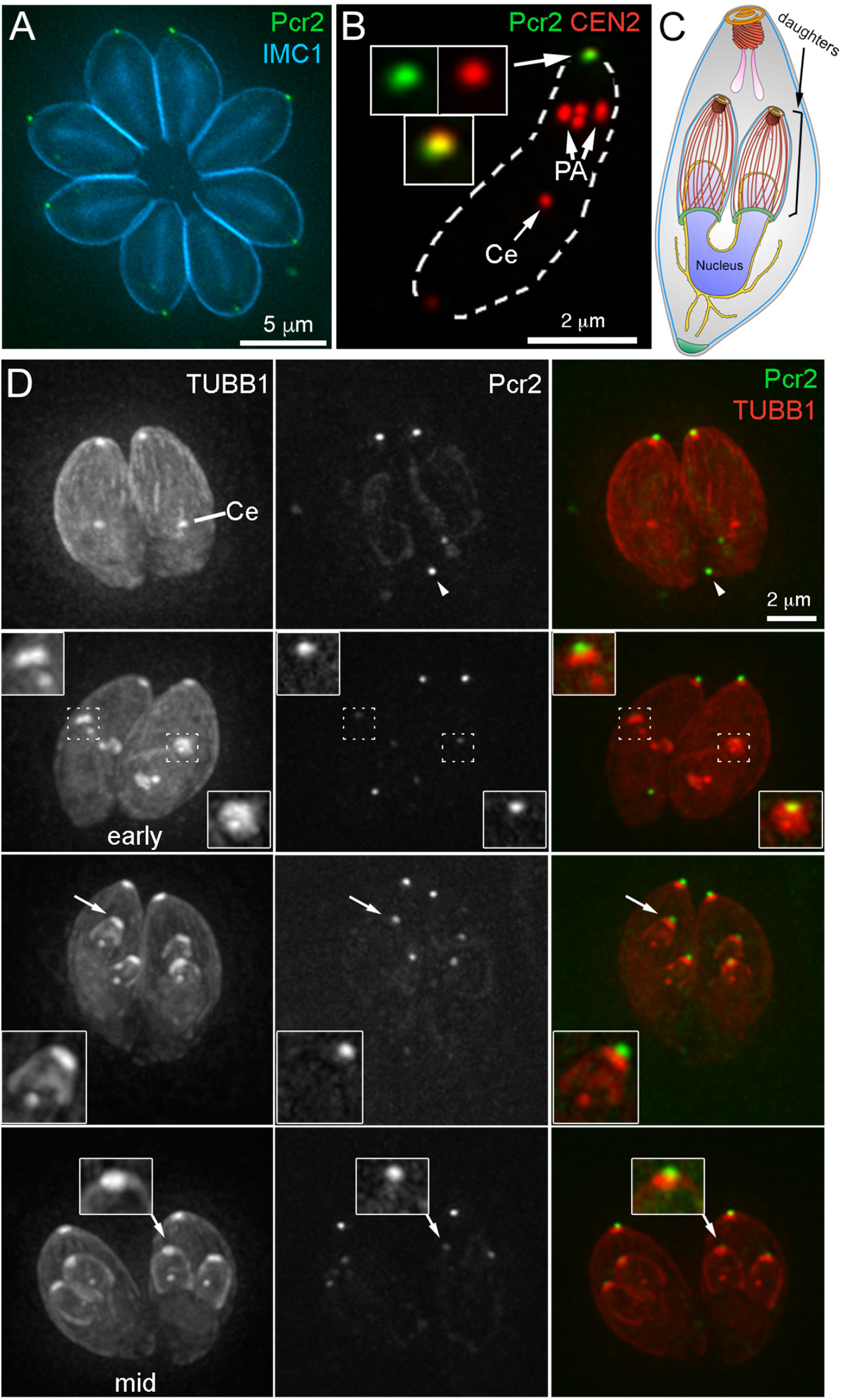
Pcr2 is recruited to the preconoidal region at an early stage of daughter formation. **A**. Projection of deconvolved wide-field images of intracellular *Pcr2-mNeonGreen* 3’ endogenous tag parasites (green) labeled with a mouse anti-IMC1 antibody and a secondary goat anti-mouse Alexa350 antibody (cyan). **B**. Projection of deconvolved wide-field images of TX-100 extracted extracellular *Pcr2-mNeonGreen* 3’ endogenous tag parasites (green) labeled with rat anti-CEN2 antibody and a secondary goat anti-rat Cy3 antibody (red). Insets (shown at 2X, contrast enhanced to display weaker signals) include the preconoidal region (arrow). Ce: centrioles. PA: peripheral annuli. **C**. Drawing depicting a dividing parasite with daughters developing inside the mother cell. For simplification, the cortical microtubules of the mother parasite are not shown. **D**. Montage showing projections of deconvolved wide-field images of intracellular *Pcr2-mNeonGreen* 3’ endogenous tag parasites transiently expressing mAppleFP-*β*1-tubulin (TUBB1, red) from a *T. gondii* tubulin promoter. The cortical microtubules in the mother parasites are present and clearly seen in the single slices of the 3-D stack, but not clearly visible in these projections due to the decreased contrast in the maximum intensity projection for weaker signals. *Pcr2* (green) is recruited to the newly formed apical cytoskeleton as soon as the daughters are detectable. Top row: interphase parasites. Row 2-4: parasites with daughters from early to mid-stage of assembly. Insets (shown at 2X, contrast enhanced to display weaker signals) include the apical region of one of the daughter parasites indicated by arrows. Ce: centrioles. Arrowhead: a Pcr2-mNeonGreen concentration is occasionally seen in the basal region of these parasites.

### Pcr2 is important for the parasite lytic cycle and invasion, but its loss does not affect apical complex structure nor parasite replication

In the *mEmeraldFP-Pcr2* knock-in parasites, the endogenous *Pcr2* gene in the parental (RHΔ*hx*Δ*ku80*) parasite has been replaced with a LoxP flanked cassette that includes the mEmeraldFP-Pcr2 coding sequence and a selectable marker, HXGPRT (Figure 4A, left panel). As described previously [7, 59-61], this arrangement was designed to allow for Cre recombinase-mediated excision of the region between the LoxP sites and thus generate a knockout mutant of the target gene. After transient expression of Cre recombinase, viable knockout mutant (*Δpcr2*) clones were generated from the *mEmeraldFP-Pcr2* knock-in parasites. This was drastically different from what had been observed with Centrin2, which is an essential gene (Numerous attempts of generating CEN2 knockout mutants using the same method all failed [55]).

**Figure 4.**
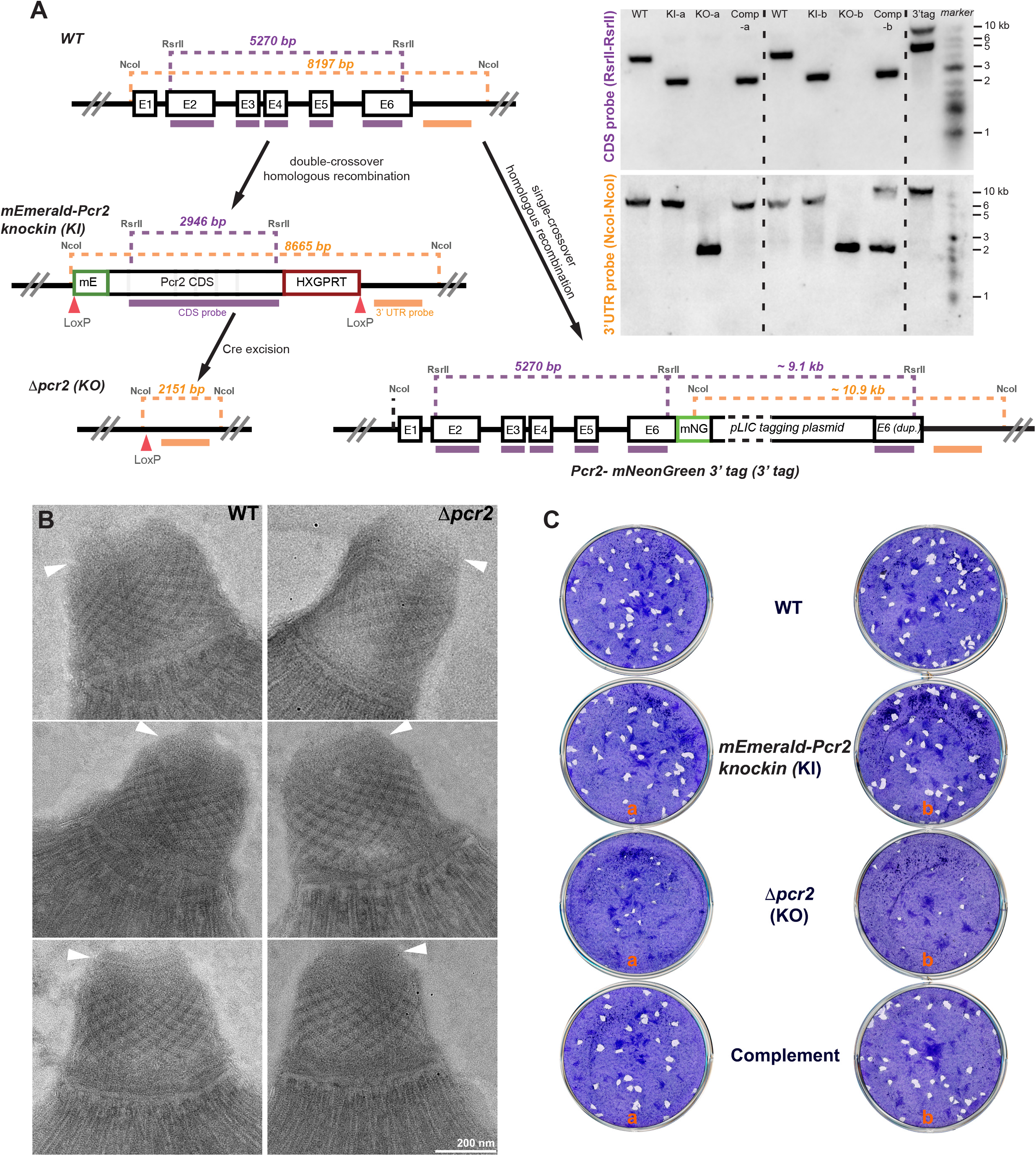
Generation of *mEmeraldFP-Pcr2* knock-in and *Δpcr2* parasites and assessment of their plaquing efficiency **A. *Left***, schematic for generating *mEmeraldFP-Pcr2* knock-in, *Δpcr2* parasites, and the *Pcr2-mNeonGreen* 3’ endogenous tagged line, and Southern blotting strategy. The positions of the restriction sites, CDS probe (purple bar) and the probe annealing downstream of the *pcr2* coding sequence (“3’ UTR probe”, orange bar) used in Southern blotting analysis and the corresponding DNA fragment sizes expected are shown (also see Materials and Methods for expected hybridization patterns). ***Right***, Southern blotting analysis of the *pcr2* locus in RH*ΔhxΔku80* (WT), *mEmeraldFP-Pcr2* knock-in (KI), *Δpcr2* (KO), complemented (Comp) parasites and the *Pcr2-mNeonGreen* 3’ tagged line (“3’tag”). The box representing the pLIC tagging plasmid is not drawn to scale. Two sets (a and b) of independently generated knock-in, knockout, and complemented lines were analyzed. **B**. EM examination of the apical complex in the RH*ΔhxΔku80* parental (WT, left column) and *Δpcr2* (right column) parasites that had been incubated with the calcium ionophore A23187 (which induces conoid protrusion), followed by TX-100 treatment. Arrowheads: preconoidal rings. **C**. Plaques formed by RH*ΔhxΔku80* (WT), *mEmeraldFP-Pcr2* knock-in (KI), *Δpcr2* (KO), and complemented (Complement) parasite lines. Plaque assays for two independent sets of knock-in, knockout and complemented lines are shown. Nine days after inoculation, the cultures were fixed with 70% ethanol and then stained with crystal violet. “plaques” are cleared spaces where the HFF monolayers were destroyed by recurring cycles of parasite invasion, replication and egress.

*Δpcr2* clones were identified based on loss of mEmeraldFP-Pcr2 fluorescence, and confirmed with genomic PCR (data not shown). Two sets (a and b) of independently generated knock-in, and knockout lines were also analyzed by Southern blotting (Figure 4A, right panel), which proved that the *pcr2* loci in parental, *mEmeraldFP-Pcr2* knock-in, and *Δpcr2* parasites, are as predicted. Complemented parasite lines were generated from *Δpcr2* parasites by reintroducing the knock-in plasmid. The Southern blot band pattern suggests that the plasmid was integrated into different locations of the genome in the two complemented lines, one of which (a) was the *pcr2* locus.

We examined the *Δpcr2* parasites using electron microscopy (EM) to determine the impact of loss of Pcr2 on the structure of the conoid. The parasites were incubated with the calcium ionophore A23187 (which increases intra-parasite [Ca^2+^] and induces conoid protrusion), followed by TX-100 treatment. We found that the loss of Pcr2 does not cause a major change in the morphology of the protruded conoid (Figure 4B). Our visual impression is that the staining pattern of the preconoidal rings in the *Δpcr2* parasites sometimes appears to be less defined than in the wild-type parasites, although the significance of this observation is unclear. The preconoidal rings are consistently present in the *Δpcr2* parasites (12 out of 13 parasites imaged). Consistent with the lack of gross structural defects, ectopically expressed Pcr1 and Pcr3 are found at the apex of the *Δpcr2* parasites (Figure S3), similar to what was observed in wild-type parasites (Figure 2).

Despite the lack of overt structural impact, the loss of Pcr2 does have a strong effect on the lytic cycle. When the parental (RH*ΔhxΔku80*), *mEmeraldFP-Pcr2* knock-in, *Δpcr2*, or complemented parasites were allowed to infect confluent cultures of human foreskin fibroblasts (HFF) (Figure 4C), the *Δpcr2* parasites formed significantly fewer and smaller plaques (Table 1). The cytolytic efficiency of the *Δpcr2* parasites is restored by complementation. The corresponding lines in sets *a* and *b* of knock-in, knockout, and complemented parasites behave similarly.

**Table 1.** Quantification of plaque assay for the four *T. gondii* strains. s.d.: Standard Deviation. Cytolytic efficiency (CE) = (Total area of the host cell monolayer lysed by a parasite line)/(Total area lysed by the WT parasite). P-values from unpaired Student’s t-tests are indicated on the right. P_n_: P-values for comparison of number of plaques. P_a_: P-values for comparison of average area of plaques.

| Strain | Number of plaques± s.d. | average area (mm<sup>2</sup>)± s.d. | CE (%WT) | *RHΔhxΔku80* (WT) | *mEmeraldFP-Pcr2 knock-in* | *Δpcr2* | complement |
| --- | --- | --- | --- | --- | --- | --- | --- |
| *RHΔhxΔku80* (WT) | 41±3 | 0.96±0.15 | 100 |  | p<sub>n</sub> = **0.039**<br>p<sub>a</sub> = 0.747 | p<sub>n</sub> = **0.0002**<br>p<sub>a</sub> = **0.015** | p<sub>n</sub> = 0.239<br>p<sub>a</sub> = 0.649 |
| *mEmeraldFP-Pcr2 knock-in* | 35±2 | 1.02±0.24 | 89.7 |  |  | p<sub>n</sub> = **0.0008**<br>p<sub>a</sub> = **0.034** | p<sub>n</sub> = 0.658<br>p<sub>a</sub> = 0.869 |
| *Δpcr2* | 9±3 | 0.31±0.03 | 7.5 |  |  |  | p<sub>n</sub> = **0.049**<br>p<sub>a</sub> = **0.046** |
| complement | 32±10 | 1.05±0.29 | 89.1 |  |  |  |  |

To determine specifically how Pcr2 knockout results in significantly reduced host cell destruction, we examined its impact on invasion, replication, and egress, the three components of the lytic cycle. By counting the number of parasites per vacuole at 12, 24, and 36 hours after infection, we found that the parental (RH*ΔhxΔku80*), *mEmeraldFP-Pcr2* knock-in, *Δpcr2*, or complemented parasites all replicate at a similar rate. Therefore, the loss of Pcr2 does not have a significant impact on parasite replication (Figure 5A).

**Figure 5.**
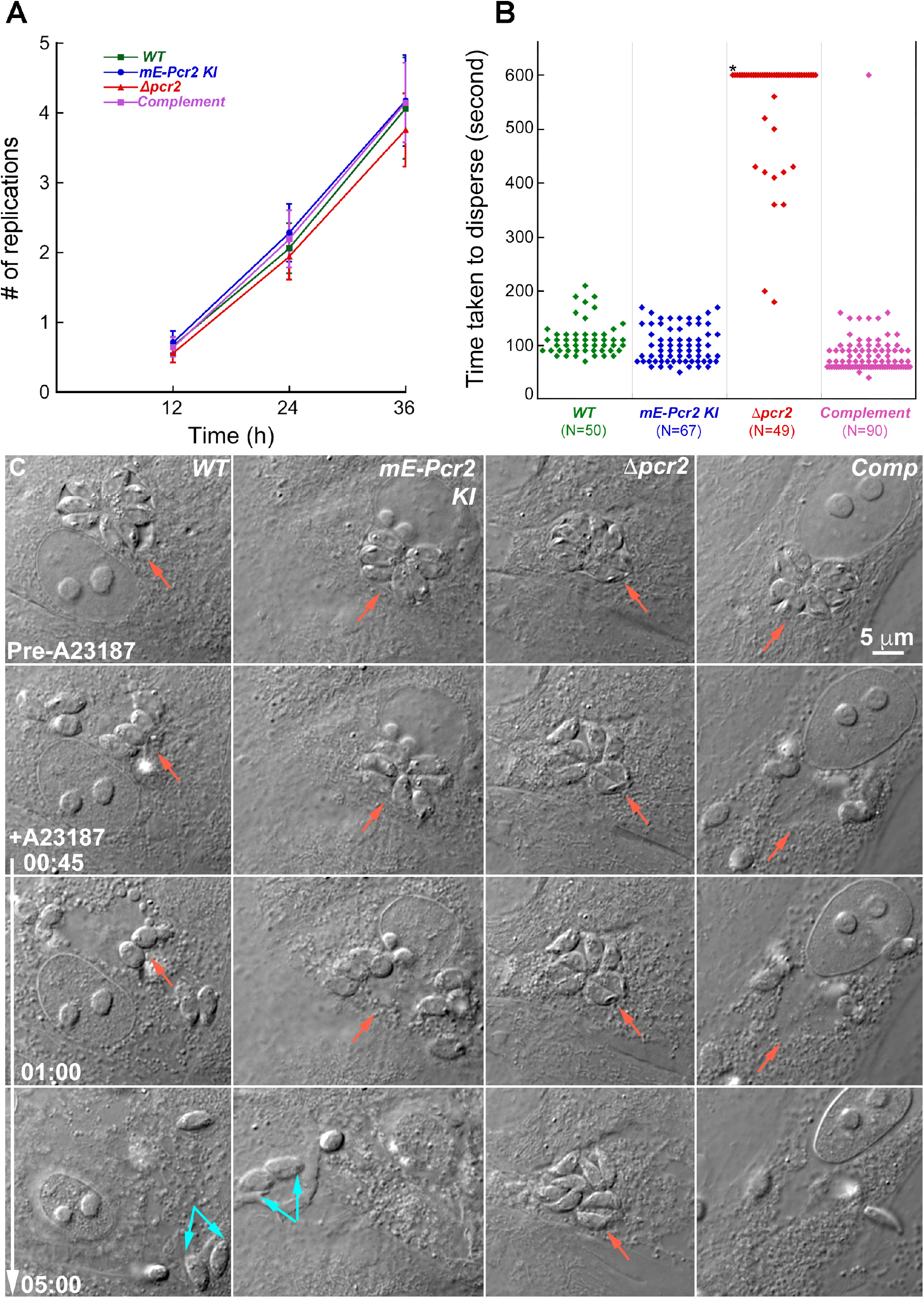
Parasite replication is not affected, but egress is impaired in *Δpcr2* parasites. **A**. The average number of replications at 12, 24 or 36 hrs after infection in four independent experiments for RH*ΔhxΔku80* (*WT*), *mEmeraldFP-Pcr2* knock-in (*mE-Pcr2 KI*), knockout (*Δpcr2*), and complemented (Comp) parasites. Error bars: standard error **B**. Dot plots of time taken to disperse after treated with 5 µM A23187 for intracellular RH*ΔhxΔku80* (*WT*), *mEmeraldFP-Pcr2* knock-in (*mE-Pcr2 KI*), knockout (*Δpcr2*), and complemented parasites. N: total number of vacuoles analyzed in 4 experiments, in which the parasite egress was monitored by time-lapse microscopy with 10-second intervals. *: *Δpcr2* parasites in 37 out of 49 vacuoles failed to disperse from the parasitophorous vacuole during the 600 sec observation period. **C**. Images selected from time-lapse experiments of intracellular WT, *mE-Pcr2* KI, *Δpcr2*, and complemented parasites treated with 5 µM A23187 (also see Video S2). A23187 was added immediately before the beginning of the time-lapse. Red arrows indicate the positions of the parasitophorous vacuole in each image. Cyan arrows indicate some of the egressed parasites that have invaded into a new host cell.

### Loss of Pcr2 results in significant defects in parasite invasion

To assess parasite invasion, we used an assay in which invaded (i.e., intracellular) and non-invaded (i.e., extracellular) parasites are distinguished based on accessibility of the parasite surface, before and after permeabilizing cells, to an antibody against a major <u>s</u>urface <u>a</u>nti<u>g</u>en (SAG1) of the parasite [22, 62]. Consistent with the smaller number of plaques formed in the plaque assay, the invasion efficiency of *Δpcr2* parasites is significantly lower than the parental lines (Table 2). The invasion efficiency was restored in the complemented line.

**Table 2.** Quantification of invasion for the four *T. gondii* strains. The number of intracellular parasites per field was counted in ten fields per strain, in each of three independent biological replicates. s.e.: Standard error of the mean. P-values from unpaired Student’s t-tests are indicated on the right.

| Strain | Number invaded ± s.e. | % WT | *RHΔhxΔku80* (WT) | *mEmeraldFP-Pcr2 knock-in* | *Δpcr2* | complement |
| --- | --- | --- | --- | --- | --- | --- |
| *RHΔhxΔku80* (WT) | 45.4±6.0 | 100 |  | p = 0.78 | p = **0.049** | p = 0.33 |
| *mEmeraldFP-Pcr2 knock-in* | 43.1±5.0 | 96 |  |  | p = **0.038** | p = 0.18 |
| *Δpcr2* | 23±2.5 | 51 |  |  |  | p = **0.005** |
| complement | 53.6±3.9 | 120 |  |  |  |  |

### Loss of Pcr2 results in significant defects in parasite egress

In contrast to the rapid dispersal of wild-type parasites after egress, in the *Δpcr2* cultures, clusters of parasites are often observed in the vicinity of a lysed vacuole, suggesting that these parasites do not egress efficiently. To examine parasite behavior during egress, we carried out time-lapse microscopy to monitor egress induced by calcium ionophore (A23187) (Figure 5B-C, Video S2). Unlike wild-type parasites, *Δpcr2* parasites in most vacuoles (37 out of 49 vacuoles) failed to disperse even 10 min after A23187 treatment (Figure 5B). For the parental, knock-in and complemented lines, it is common to observe reinvasion immediately after egress (Figure 5C, cyan arrows). In contrast, reinvasion of egressed *Δpcr2* parasites was never observed in a substantially longer recording period (> 10 min), consistent with the low invasion efficiency demonstrated by the immunofluorescence-based assay (Table 2).

### Δpcr2 parasites can secrete effectors to lyse the host cell during calcium-induced egress and the localization and secretion of the major adhesin, MIC2, are normal

Lysis of the host cell in response to elevated calcium [38] occurs normally in *Δpcr2* parasites. Shortly after the *Δpcr2* infected culture was exposed to A23187, nuclear labeling by a cell-impermeant DNA binding dye (DAPI) included in the culture medium was detected within the now-permeable host cell. DAPI in the culture medium and in the cytoplasm is not seen because its fluorescence is low until it binds to DNA. DAPI fluorescence first appears starting at the rim of the host cell nucleus, then spreads inwards (Figure 6A, Video S3). The host cell also showed other symptoms of lysing, including the roundup of mitochondria, blebbing and contraction (Videos S2-S3). Note that the dynamics of the DAPI binding captures the local nature of the initiation of host cell lysing, as the fluorescence always appears first on the side of the nucleus closer to the parasitophorous vacuole. The nuclei of neighboring uninfected host cells are not labeled by DAPI (Figure 6A), further confirming that the host cell lysing is a parasite driven process.

**Figure 6.**
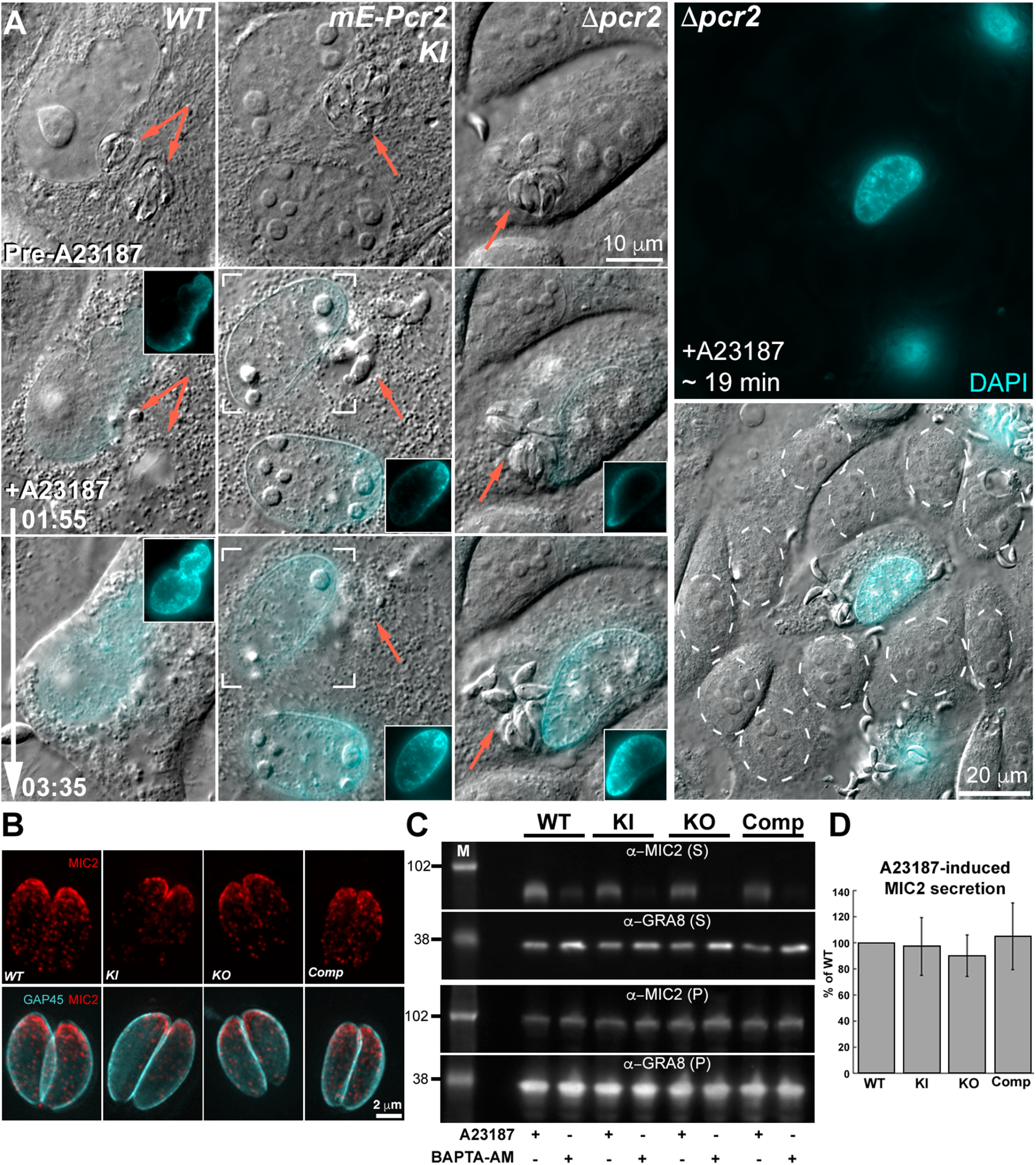
Calcium ionophore-induced micronemal secretion is not significantly affected in *Δpcr2* parasites. **A**. Images selected from time-lapse experiments of intracellular RH*ΔhxΔku80* (*WT*), *mEmeraldFP-Pcr2* knock-in (*mE-Pcr2 KI*), and knockout (*Δpcr2*) treated with 5 µM A23187 (also see Video S3). The cell-impermeant DNA-binding dye, DAPI, was added to the medium to monitor the permeabilization of the host cell. *Δpcr2* parasites are able to secrete effectors that lyse the host cell upon A23187 treatment, indicated by DAPI entering the host cell nucleus and binding to DNA, as well as by the dramatic change in the morphology of the host cell (see Videos S2-S3). Insets are DAPI images of the nuclear region of the host cell shown at 0.5X. Brackets in the *mE-Pcr2 KI* panels indicate the host cell nucleus included in the insets. Contrast was adjusted so that the DAPI labeling at the rim of the nucleus is easily visible. The nuclei of uninfected fibroblasts (marked by dashed circles) remained unlabeled by DAPI ∼19 min after A23187 treatment as shown in the larger field of view images in the right-hand column. **B**. Projections of deconvolved wide-field fluorescence images of intracellular WT, *mE-Pcr2* KI, *Δpcr2*, and complemented (Comp) parasites labeled with a mouse anti-MIC2 (red), a rat anti-GAP45 (cyan) and corresponding secondary antibodies. **C**. Western blots of the secreted (supernatant, S) and unsecreted (pellet, P) fractions of WT, *mE-Pcr2* KI, *Δpcr2*, and complemented (Comp) parasites after A23187 or BAPTA-AM (a calcium chelator; negative control) treatment. The blots were probed by antibodies against MIC2 and GRA8. M: molecular weight markers, the masses of which are indicated in kDa by the numbers on the left. **D**. Levels of MIC2 in the secreted fractions relative to that from the wild-type in 3 independent biological replicates. For each sample, the MIC2 secretion upon A23187 stimulation is normalized against GRA8 in the pellet from the same sample. Error bars: standard error.

Efficient host cell lysing suggests functional secretion of micronemal lytic proteins by the parasite. To further confirm this hypothesis, we investigated the impact of Pcr2 knockout on microneme distribution and secretion directly by examining the intracellular distribution and A23187-induced secretion of <u>Mic</u>ronemal Protein <u>2</u> (MIC2), a major transmembrane adhesin important for parasite motility [44, 45, 63]. Immunofluorescence analysis showed no significant differences in MIC2 distribution between intracellular wild-type and *Δpcr2* parasites (Figure 6B). A23187-induced MIC2 secretion from the *Δpcr2* parasites is also not significantly different from the wild-type, knock-in and the complemented lines (Figure 6C-D). This indicates that the major egress defect seen in the *Δpcr2* parasite is not due to the loss of adhesin secretion. For the secretion assays, the dense granule protein GRA8 was used as a control, the secretion of which is negatively regulated by calcium as previously reported for other dense granule proteins [64].

### Δpcr2 parasites move spasmodically

Close examination of the behavior of the *Δpcr2* parasites revealed that these parasites do become motile shortly after treatment with 5 *μ*M A23187 (Video S2-S3). However, unlike the parental and complemented parasites, whose movements led to nearly synchronous dispersal, *Δpcr2* parasites moved spasmodically and displayed poor efficiency for egress. Eventually some parasites manage to escape after vacillating multiple times before leaving the lysed vacuole. Previously, we discovered that another component of the apical complex, the <u>A</u>pical complex Lysine(<u>K</u>) <u>m</u>ethyltransferase (AKMT), is an activator for parasite motility [7]. Knockout of AKMT leads to drastically impaired invasion and egress. Comparison between the *Δpcr2* and *Δakmt* parasites revealed that these two mutants behave quite differently (Figures 7-8). Both mutants respond to the A23187 treatment by lysing the host cells (Figure 8A, Video S4), but their behaviors in motility are different. While the *Δakmt* parasites were in almost total paralysis, the *Δpcr2* parasites could still move but in a fitful manner. This indicates that these two apical proteins contribute differently to parasite motility. The type of spasmodic motion observed in the *Δpcr2* parasites has not been reported in other previously characterized motility mutants [7, 12, 13, 41-43, 46].

**Figure 7.**
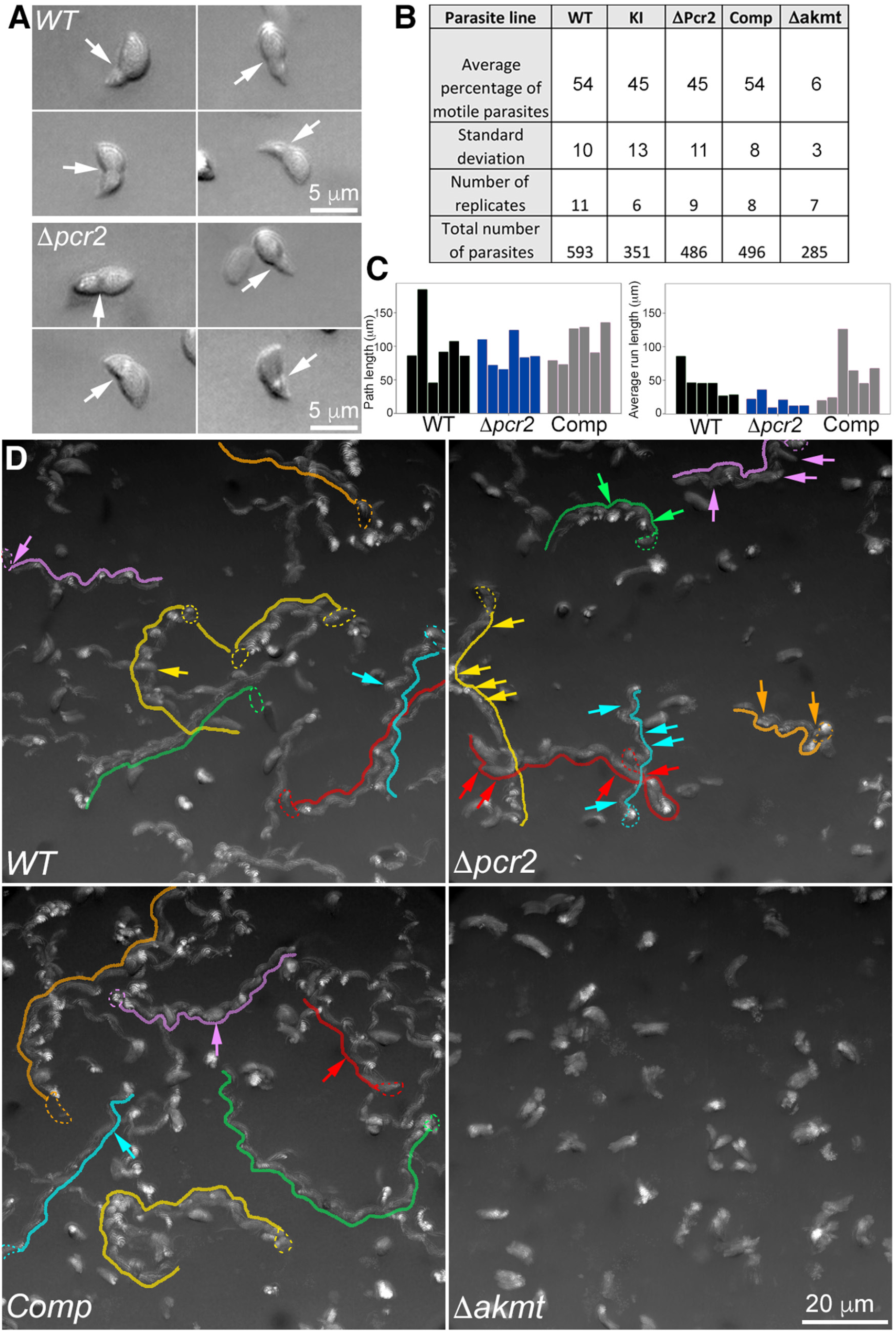
The *Δpcr2* parasite moves spasmodically in Matrigel. **A**. Four examples each of wild-type and *Δpcr2* parasites that displayed constriction during movement in 50% Matrigel. **B**. Table showing percentage of motile RH*ΔhxΔku80* (*WT), mE-Pcr2* Knock-in (KI), *Δpcr2*, complemented (Comp), and *Δakmt* parasites in the 3-D motility assay. **C**. Bar graphs that show the path length and average run length of the trajectories highlighted in D. Average run length is calculated by dividing the path length by the number of runs within each trajectory. The number of runs is defined as 1+ number of sustained (> ∼11 sec) pauses. **D**. Movement tracks for WT, *Δpcr2*, complemented (Comp), and *Δakmt* parasites generated by projections of 3-D motility timelapses. See Videos S6-S7. To make these 2-D images of parasites’ paths through the 3-D gel, a projection of the 3-D stack at each time point was first generated by Stack focuser (ImageJ/Fiji), then 150 consecutive timeframes in the projected sequence were compressed into a single frame. Six traces each for WT, *Δpcr2*, and complemented parasites are highlighted by traces drawn by hand. Some of the pauses in the parasite movement that lasted for 7 frames (∼ 11 sec) or longer in these traces are indicated with arrows of the same color.

**Figure 8.**
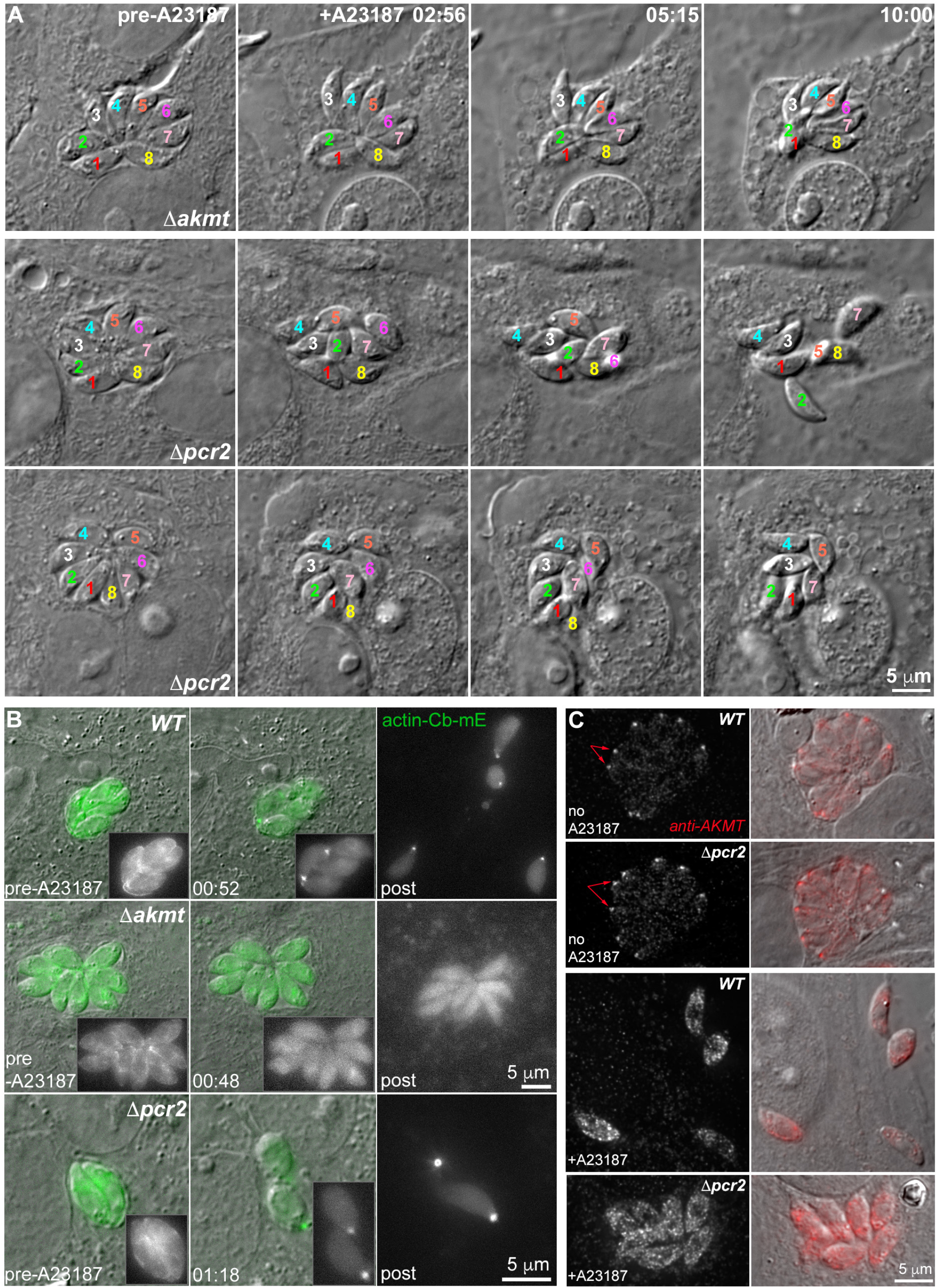
Pcr2 functions differently from another motility regulator, AKMT, and Pcr2 knockout does not block actomyosin activity. **A**. Images selected from time-lapse experiments of intracellular *Δakmt* and *Δpcr2* parasites treated with 5 µM A23187. Note that *Δakmt* parasites, which are largely immobile, maintained their organization within the vacuole after lysis of the host cell. However the position and orientation of the *Δpcr2* parasites shifted during the time lapse due to sporadic parasite movement (see Video S4). **B**. Actin-chromobody-mEmerald (actin-Cb-mE) distribution before and after A23187 treatment in RH*ΔhxΔku80* parental (*WT*), *Δakmt* and *Δpcr2* parasites. The grayscale actin-Cb-mE fluorescence images are projections of the image stack at the corresponding time point. **C**. Localization of AKMT in the RH*ΔhxΔku80* parental (*WT*) and *Δpcr2* parasites before and after ∼ 5 min 5 µM A23187 treatment. Before exposure to A23187, AKMT is concentrated at the apical end (arrows) of intracellular WT and *Δpcr2* parasites. The increase in intra-parasite [Ca^2+^] caused by A23187 treatment triggers the dispersal of AKMT from the parasite apical end in both the parental and the *Δpcr2* parasites. AKMT was labeled by immunofluorescence using a rat anti-AKMT antibody and a secondary goat anti-rat Cy3 antibody. The grayscale anti-AKMT fluorescence images are projections of deconvolved image stacks.

The spasmodic motion of the *Δpcr2* parasite during egress is characterized by sudden forward movement followed by long pauses or backward sliding. This could be due to an inherent defect in maintaining directional parasite movement. Alternatively, it could be caused by a failure in overcoming resistance from the surrounding host subcellular structures. To examine the parasite motile behavior in a host-cell-free environment, we used a 3-D motility assay previously developed by the Ward Lab [65], in which the parasites are allowed to move in a 3D gel (Matrigel) that approximates the extracellular matrix of mammalian tissue. In the original form of the assay, parasite position was determined by nuclear tracking, enabled by nuclear labeling with the cell-permeant DNA dye Hoechst 33342 and fluorescence imaging with Ultraviolet (UV) excitation. We chose to use label-free Differential Interference Contrast (DIC) imaging to eliminate UV exposure to the parasite, and to enable imaging of any morphological changes the parasite undergoes while moving in the 3-D environment. Using a silicone immersion objective, for which the refractive index of the immersion oil (n=1.405) is close to that of the Matrigel (n∼1.34), high-quality DIC images can be acquired throughout a region exceeding 100 microns in depth. As reported previously [65], motile wild-type parasites move along a helical path in the Matrigel (Figure 7). Often the movement continues over tens of microns, many body-lengths of the parasite. Interestingly, constrictions are observed around some moving parasites, possibly a result of the parasite pushing through small pores in the matrix (Figure 7A, Video S5). This is similar to what is seen during invasion of a host cell, and further validates the 3-D motility assay as a physiologically relevant analysis for *Toxoplasma*. As expected, the majority of *Δakmt* parasites are immotile (Figure 7B), consistent with the paralyzed behavior observed in the induced egress assay. In contrast to the nearly immotile *Δakmt* parasites, the proportion of motile *Δpcr2* parasites was close to that of the wild-type parasites (Figure 7B). However, the movement of the *Δpcr2* parasite is noticeably less sustained (Videos S6-S7) with more frequent long pauses (Figure 7C-D, indicated by arrows in D), consistent with what was observed in the egress assay.

### Loss of Pcr2 does not block actomyosin-based motion nor AKMT dynamics

*Toxoplasma* motility is generally thought to be driven by actin polymerization and associated myosin motors [2, 6, 7, 12, 39-43]. To investigate the dynamics of actin-containing structures, several groups have used an actin-chromobody tagged with a fluorescent protein (actin-Cb-mE or actin-Cb-GFPTy) and found that the distribution of the tagged actin-Cb is sensitive to changes in intra-parasite calcium concentration, induced by BIPPO (a cGMP phosphodiesterase inhibitor) or the calcium ionophore, A23187 [13, 48, 66, 67]. The redistribution of actin-Cb in response to elevated calcium appears to be dependent on actin polymerization and myosin activity [13, 48]. Indeed, in live egress experiments, we observed that A23187 treatment induced an actin-Cb-mE accumulation at the basal end in 100% of the wild-type parasites (n= 127) (Figure 8B). In contrast, the actin-Cb-mE accumulation was not observed in ∼ 58% of the *Δakmt* parasites (n=125) when treated with A23187. In the ∼42% of A23187 treated *Δakmt* parasites that showed some basal concentration of actin-Cb-mE, the accumulation typically was not nearly as pronounced as observed in wild-type parasites. This is largely in agreement with a prior observation that BIPPO for the most part failed to induce actin-Cb-GFPTy basal accumulation when AKMT was knocked down [13]. We previously discovered that jasplakinolide, which stabilizes actin filaments, compensates the defect in magnitude of the cortical force generated by *Δakmt* parasites in laser trap measurements [68]. Thus, both lines of evidence suggest that AKMT might regulate parasite motility by controlling actin polymerization. While the *Δpcr2* parasites also have a pronounced motility defect, the basal accumulation of actin-Cb-mE occurred in 95% of these parasites (n= 159) when stimulated by A23187. This raises the possibility that Pcr2 acts downstream of or in parallel with the actomyosin machinery and AKMT. Consistent with this idea, we found that the loss of Pcr2 does not affect the apical localization of AKMT, and that the calcium-triggered dispersal of AKMT from the parasite apex [7] still occurs in the *Δpcr2* parasites (Figure 8C).

### Spasmodic movement contributes to the invasion defect of Δpcr2 parasites

Parasite motility is required for both egress and invasion. Many motility-relevant genes are involved in both processes. On the other hand, invasion does have its unique structural requirements [14, 15, 17, 18, 22]. We were therefore curious to see whether an invading *Δpcr2* parasite displays a similar type of spasmodic motility as observed during egress and during movement in the 3-D matrix, and if so, how the fitful nature of the movement might affect the efficiency of invasion. To address this, we carried out live-cell imaging and specifically examined the action of parasites after they had initiated contact with a host cell. In these experiments, freshly harvested extracellular parasites were settled down onto the host cell monolayer by slow speed centrifugation or by a pre-incubation on ice. After parasites had settled down, the cultures were then imaged by time-lapse DIC microscopy at 37°C (Figure 9, Video S8). We then counted the number of invasion or attempted invasion events recorded in the videos for four parasite lines: wild-type, Pcr2-knock-in, *Δpcr2*, and complemented. We observed 264 events for wild-type, 194 for knock-in, 27 for *Δpcr2*, and 174 for complemented parasites. Of these, > 99% (i.e. 263 of 264) of the wild-type, ∼ 98% of the knock-in and ∼ 97% of the complemented parasites completed the invasion. In contrast, ∼ 63% of the invasion attempts (17 of 27) by *Δpcr2* parasites stalled or were aborted after the parasite managed a partial entry (Figure 9D-E, Video S8). Among the 27 *Δpcr2* parasites analyzed, constrictions were observed in 12 and not found in 10 parasites. For the remaining 5 parasites, the formation of constriction could not be determined definitively. In comparison, constrictions were consistently observed in invading parasites of the Pcr2-expressing strains (31 out of 33 randomly selected wild-type parasites, 25 out of 31 Pcr2 knock-in parasites, 26 out of 28 complemented parasites).

**Figure 9.**
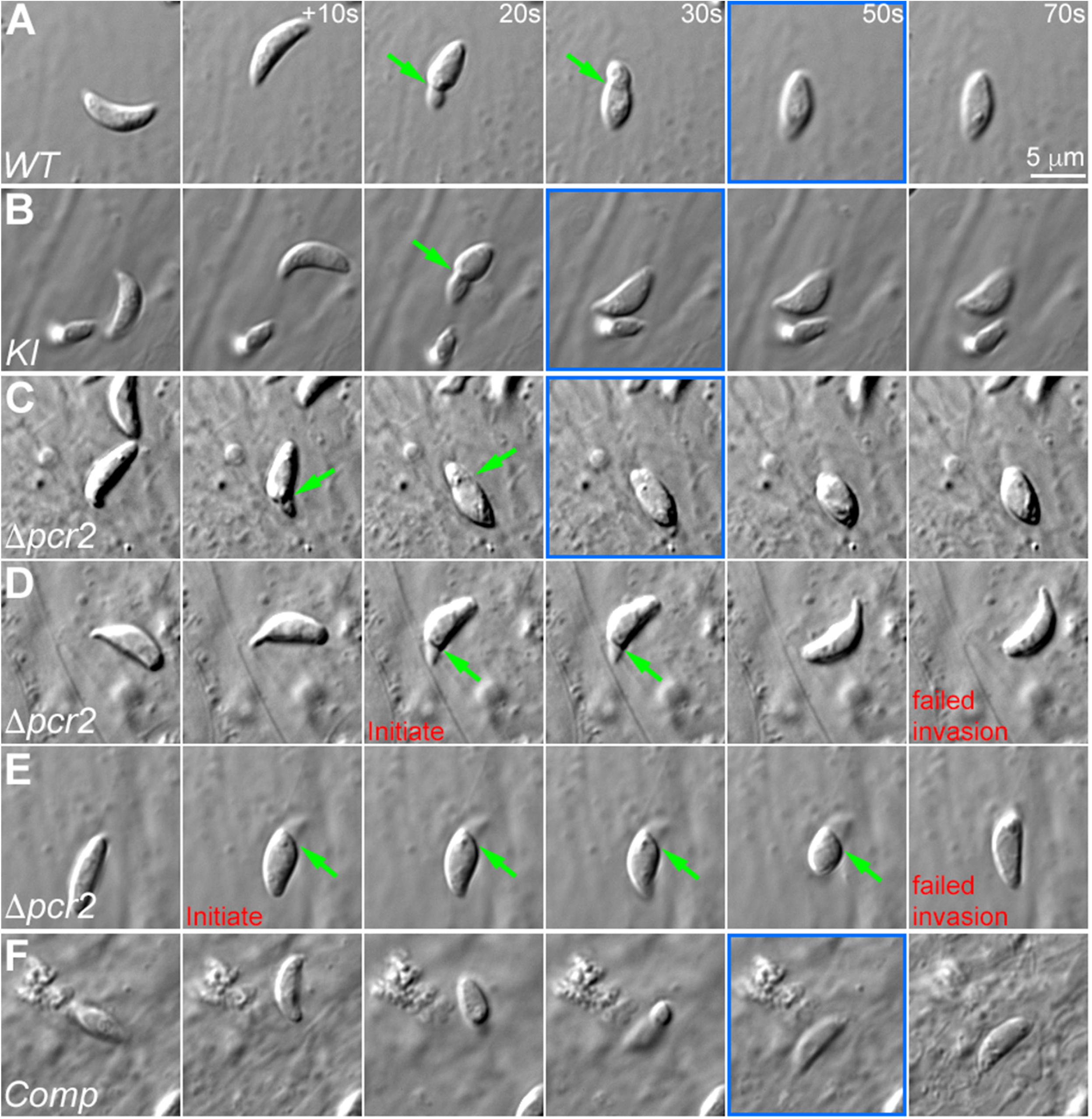
The *Δpcr2* parasite moves fitfully during invasion. Images selected from time-lapse recording of RH*ΔhxΔku80* (*WT*, **A**), *mEmeraldFP-Pcr2* knock-in (*KI*, **B**), knockout (*Δpcr2*, **C-E**), and complemented (*Comp*, **F**) parasites in the process of invasion or attempted invasion. D and E show two examples of abortive invasion by the *Δpcr2* parasite. The frames where the invasion has completed are marked in blue. Green arrows: constrictions formed during invasion. Also see Video S8.

Previous studies have shown that a ring-like moving junction forms at the site of the constriction that develops during normal parasite invasion [14-21]. To determine whether the assembly of this moving junction occurs normally in the *Δpcr2* parasites, we examined the localization of RON4 (Rhoptry Neck Protein 4), which is found at the neck region of rhoptries in intracellular parasites and is recruited to the moving junction during parasite invasion [14, 15]. In quiescent intracellular parasites, there is no detectable difference in RON4 localization between knock-in and *Δpcr2* parasites (Figure 10A). To examine RON4 localization in invading parasites, we incubated the parasites with host cells for a short period of time, fixed the cultures, incubated with a rabbit anti-SAG1 antibody (to label the part of the parasite plasma membrane not yet shielded by the parasitophorous vacuole), then detergent-permeabilized and incubated with a mouse anti-RON4 (to highlight the moving junction) plus a rat anti-GAP45 antibody (to label the Inner Membrane Complex (IMC) of the entire parasite) (Figure 10B). Parasites in these labeled cultures that had developed a constriction were first identified by DIC, and then imaged by fluorescence in the SAG1, RON4, and GAP45 channels. Four distinct patterns were observed: 1) SAG1 labeling predominantly on the parasite plasma membrane distal to the constriction (i.e., on the basal side) with a normal-looking RON4 ring at the constriction; 2) SAG1 labeling of the entire plasma membrane of the parasite with no detectable RON4 ring at the constriction, 3) SAG1 labeling of the entire plasma membrane of the parasite with a normal-looking RON4 ring at the constriction, 4) SAG1 labeling predominantly distal to the constriction with no detectable RON4 ring (Figure 10C-D). The latter two groups accounted for a small minority of the total and could reflect either technological limitations or true biological variation present in a very small fraction of parasites. The majority of the parasites were found in the first two groups. For the wild-type, knock-in, and complemented parasites, approximately 80% of the parasites that formed a constriction also assembled a well-defined RON4-marked moving junction at the constriction. In these parasites, the invaded portion of the parasite is well shielded by the parasitophorous vacuole and the moving junction, as indicated by the accessibility of the SAG1 antibody (before permeabilization with detergent) predominantly to only the extracellular portion of the parasite. For *Δpcr2* parasites, that labeling pattern is seen in less than 30% of the parasites that form a constriction. In contrast, in more than 50% of the constricted *Δpcr2* parasites, RON4 labeling at the constriction was not detected and SAG1 labeling was seen over the entire parasite, indicating poor sealing of the parasitophorous vacuole or failure of initiating a parasitophorous vacuole, leaving the parasite membrane on both sides of the constriction exposed, which could contribute to, or result from, compromised invasion.

**Figure 10.**
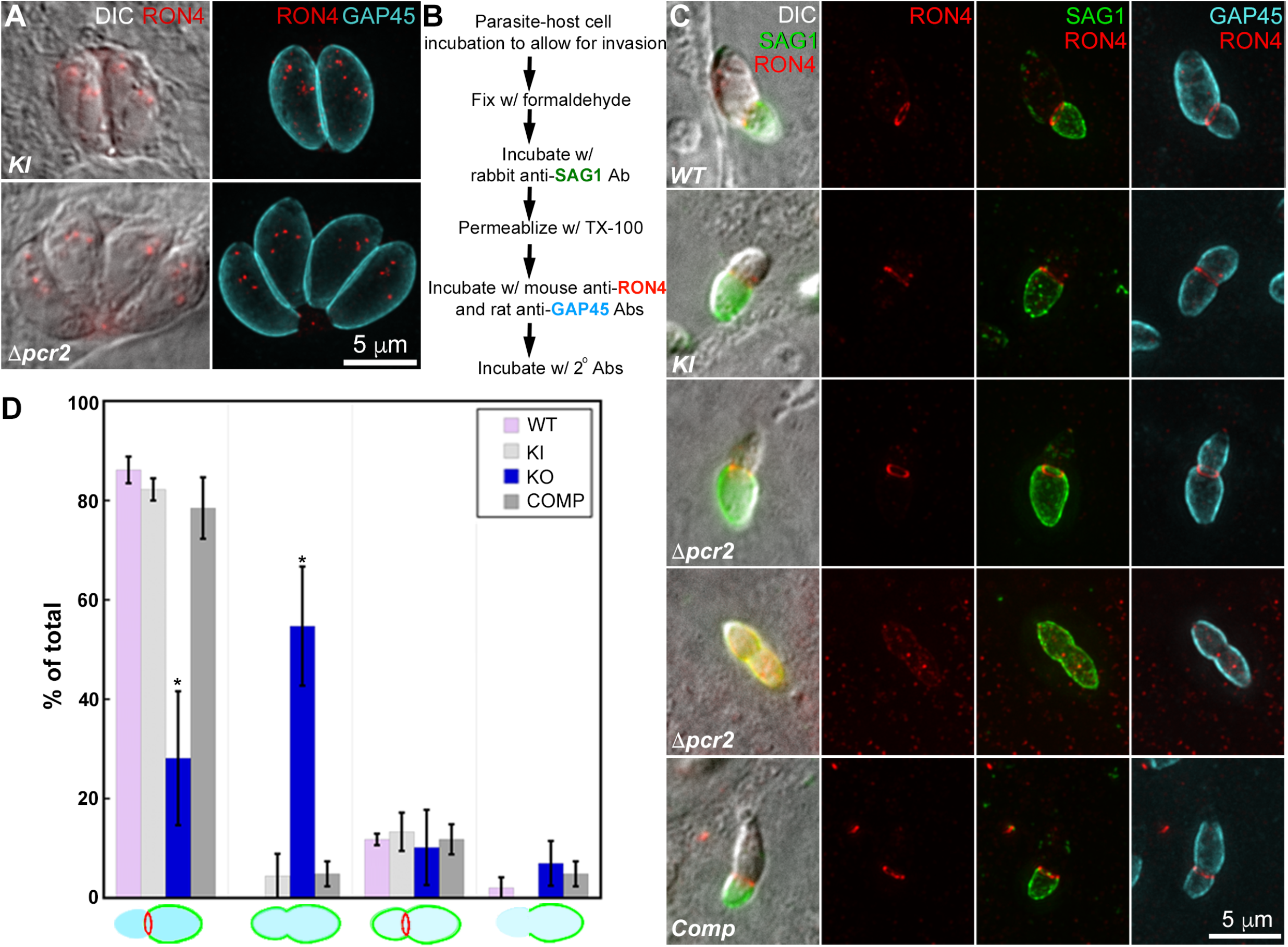
*Δpcr2* parasites are defective in assembling the moving junction. **A**. Projections of deconvolved wide-field fluorescence images of <u>intracellular</u> *mE-Pcr2* KI (*KI*) and *Δpcr2* parasites labeled with a mouse anti-RON4 (red), a rat anti-GAP45 (cyan) and corresponding secondary antibodies. **B**. Outline of a pulse invasion assay to analyze the assembly of the moving junction (marked by anti-RON4 labeling), and the differential accessibility of the intracellular vs extracellular portion of the invading parasites to antibody labeling of the SAG1 surface antigen. **C**. DIC and projections of deconvolved wide-field fluorescence images of RH*ΔhxΔku80* (*WT*), *mEmeraldFP-Pcr2* knock-in (*KI*), knockout (*Δpcr2*) and complemented (Comp) parasites, in which SAG1 (green) RON4 (red), and GAP45 (cyan) were labeled by immunofluorescence in the pulse invasion assay described in B. Two predominant patterns are included. **D**. Quantification of all four SAG1 and RON4 labeling patterns observed in *WT, KI, Δpcr2* and complemented (Comp) parasites from three independent biological replicates. Error bars: standard error. * P value <0.05 (unpaired Student’s t-tests), when compared with WT parasites.

## DISCUSSION

Apicomplexan motility is vital for host infection and parasite dissemination. To complete its life-cycle, a parasite has to navigate through complex intracellular and extracellular environments. Thus the parasite must travel through regions having widely different local mechanical properties: different combinations of permeability, pore size, elasticity, and friction, giving rise to large variations in the power output required of the parasite’s motile apparatus. How this dynamic interplay between imposed load and motor output occurs remains an unexplored area, as the outcome of manipulating the previously known motility-relevant genes has been largely binary; i.e., motile vs. immotile [7, 12, 13, 41-43, 46]. While the current model [47] provides a valuable framework connecting the functions of many proteins involved in motility (e.g. the IMC-anchored myosin motors for generating internal force, actin polymerization for providing tracks for the motor, actin-binding adapter proteins for linking the actomyosin complex to the transmembrane adhesins), it reflects the binary nature of the genetic manipulation experiments in that it describes a 2-state motility apparatus: the motor is always either “ON” or “OFF”.

Here we report a new motility phenotype, in which the parasite is motile, but the movement is intermittent and results in frustrated egress and invasion. One can imagine several possible causes for this type of behavior. For example, the defect could be primarily mechanical, such as an “engine” that is underpowered due to loss or malfunction of an important component (Figure 11). Another mechanical defect might be envisioned as a “broken transmission” (i.e. the internal engine performs normally, but the force is not transmitted effectively to the parasite outer surface to generate traction) (Figure 11). Arguing against the “wimpy motor” explanation is the observation that, at the molecular level, Pcr2 does not appear to have a strong impact on actin kinetics or actomyosin activity, as the knockout of Pcr2 does not affect the basal accumulation of actin-Cb-mE upon calcium ionophore treatment. However, it should be noted that the actin-Cb-mE-assay does not assess actomyosin activity in long-distance travel. Arguing against the “broken transmission” hypothesis is the observation of normal calcium-induced secretion of the major adhesin protein, MIC2 (part of the “transmission” in the motility apparatus) in the *Δpcr2* parasite. Of course it remains possible that other unknown adhesins or associated proteins involved in force transduction are affected by the loss of Pcr2. In either case, regardless of the protein molecules involved, the prediction is that the persistence of parasite movement would be load dependent. The motility apparatus could drive parasite movement when the load is low enough, but the movement would be stalled when the parasite encounters increased resistance. This hypothesis could be tested in the future by determining how the *Δpcr2* parasites move in different extracellular matrices with well-defined graduated stiffness and pore size.

**Figure 11.**
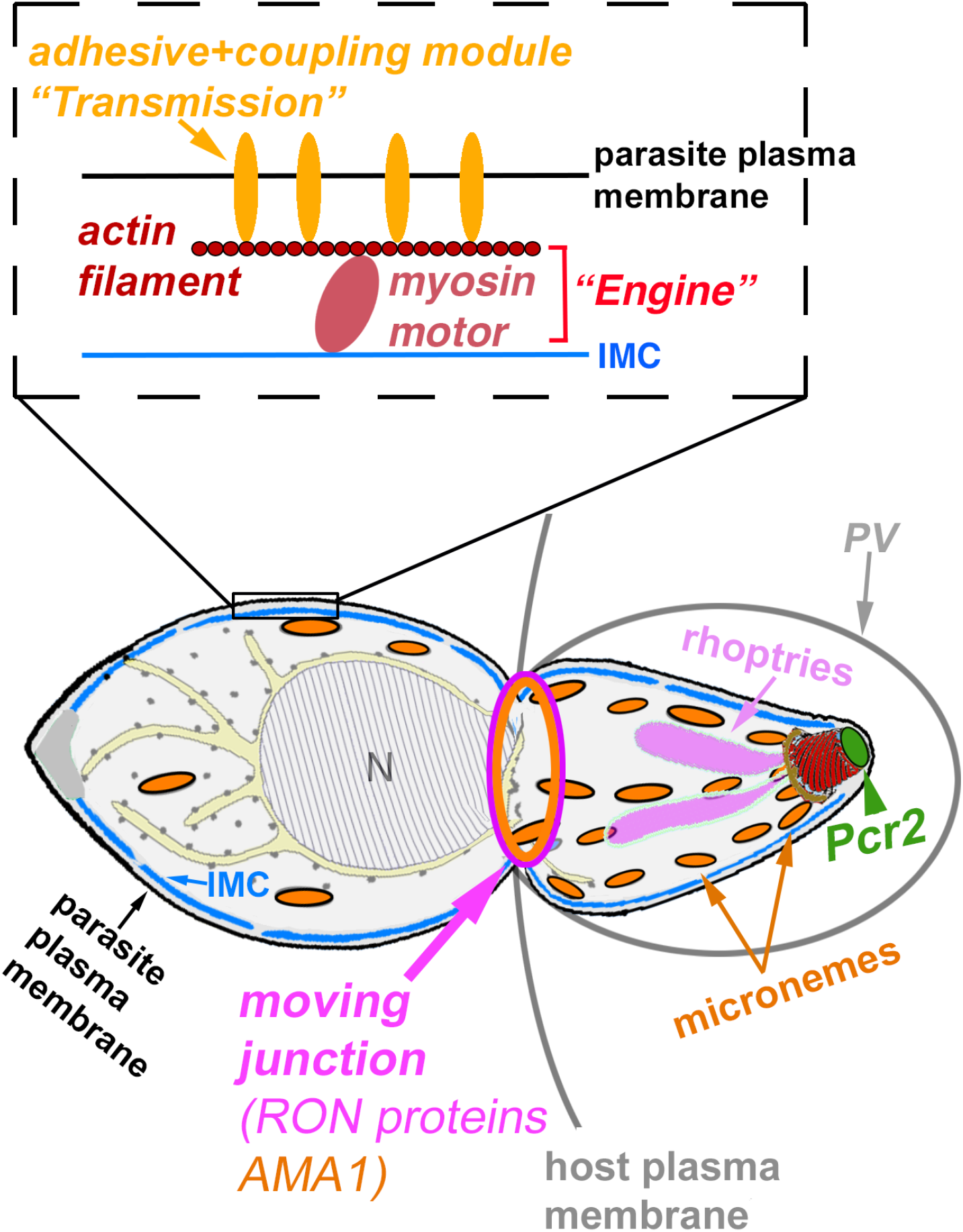
Simplified schematics of an invading parasite with the “engine” and “transmission” modules in the motility apparatus (inset) illustrated. PV: parasitophorous vacuole. N: nucleus. IMC: the Inner Membrane Complex.

Unexplained by either form of “mechanical defect” hypothesis is how Pcr2, located at the extreme apex of the parasite, would directly affect the motile apparatus, which is distributed along the parasite cortex. As an alternative to postulating a mechanical defect, the spasmodic motility could conceivably be a result of broken signaling. If sustained parasite movement requires a sustained activating signal, intermittent signaling would give rise to fitful movement. If this is the case, one would predict fitful movement of the *Δpcr2* parasite irrespective of the external load.

Aside from motility, the loss of Pcr2 also affects the assembly of the moving junction during invasion, even though Pcr2 is not detected in the moving junction in invading parasites. In our experiments, imperfect moving junction assembly, as defined by RON4 labeling, is correlated with accessibility of the entire plasma membrane of the invading parasite, indicating poor sealing of the junction or poor shielding of the parasite plasma membrane. This might be an indication of poor parasite-host-cell adhesion, hence poor traction, thus contributing to the poor invasion efficiency of the *Δpcr2* parasite. On the other hand, frustrated invasion due to stalled movement could possibly have some negative feedback on proper assembly of the moving junction. Due to the limitation of the immunofluorescence-based assay, when RON4 was not detected in the moving junction, the current work cannot distinguish whether the issue lies in the initiation or the maintenance of the RON4 ring. This needs to be directly investigated by live cell imaging in the future. It is also worth noting that, in this study, only parasites with constrictions were analyzed by immunofluorescence, because, in fixed cells, it is impossible to distinguish extracellular parasites from parasites that attempt to invade but do not form constriction. Therefore the results in Figure 10 almost certainly do not reflect the full extent of the phenotype. This is supported by the observation in live invasion experiments that some parasites that attempted to invade did not form a constriction.

The discovery of Pcr2 and the analysis of its function open new opportunities to determine how mechanical interactions with its environment impact parasite motility, and how the regulation or maintenance of persistence in parasite movement is functionally connected with and imposed onto the motility apparatus and other structures involved in invasion and egress. As Pcr2 is predominantly located at the preconoidal region, it would be interesting to determine whether Pcr2 is involved in loading other motility-relevant factors to the apical region of the parasite. Additionally, Pcr2 is a useful probe to help determine the role of the preconoidal rings as a whole in regulating parasite motility, as a signaling center for promoting persistent parasite movement, or as a mechanical facilitator for penetrating the barriers posed by the environment surrounding the parasite. Lastly, while our initial motivation of analyzing preconoidal proteins was to gain insights into the structural connection between the preconoidal rings and the conoid, the role of the preconoidal rings in organizing the conoid is still not clear and needs to be explored.

## MATERIALS AND METHODS

### T. gondii, host cell cultures, and parasite transfection

The maintenance of host cell and *T. gondii* tachyzoite parasite cultures, and parasite transfections, were carried out as previously described [59, 69, 70]. Confluent human foreskin fibroblasts (HFFs; ATCC# SCRC-1041, and HFF_hTERT; ATCC# CRL-4001) monolayers in Dulbecco’s Modified Eagle’s Medium (DMEM, VWR, 45000-316), supplemented with 1% (v/v) heat-inactivated cosmic calf serum (Hyclone, SH30087.3) and Glutamax (Life Technologies-Gibco, 35050061) were used to maintain parasite cultures. For light microscopy-based assays, HFF, African green monkey renal epithelial cells, (BS-C-1; ATCC# CCL-26), and rat aorta cells (A7r5 ; ATCC# CRL-1444) were used as host cells.

### Immunoprecipitation and Multidimensional Protein Identification Technology (MudPIT) analysis

Immunoprecipitation experiments were performed as described in [61], using *eGFP-CEN2* knock-in parasites [55] and Chromotek-GFP-Trap agarose beads (Cat#ACT-CM-GFA0050, Allele Biotechnology, CA). Protein samples were processed for MudPIT as described in [61]. MS/MS spectra were first searched using ProLuCID v. 1.3.3 [71] with a mass tolerance of 10 ppm for peptide and fragment ions. Trypsin specificity was imposed on both ends of candidate peptides during the search against a protein database containing 48080 non-redundant (NR) human proteins (combining entries from <u>GCF_000001405.40_GRCh38.p14</u> and <u>GCF_009914755.1_T2T-CHM13v2.0</u>) and 8450 *Toxoplasma gondii* NR proteins (TGGT1 entries from <u>ToxoDB-53</u>), as well as 428 common contaminants such as human keratins, IgGs and proteolytic enzymes. To estimate false discovery rates (FDR), each protein sequence was randomized (keeping the same amino acid composition and length) and the resulting “shuffled” sequences were added to the database, for a total search space of 113916 amino acid sequences. Combining all replicate runs, proteins had to be detected by at least 2 peptides and/or 2 spectral counts. Proteins that were subsets of others were removed using the parsimony option in DTASelect v. 1.9 [72] on the proteins detected after merging all runs. Proteins that were identified by the same set of peptides (including at least one peptide unique to such protein group to distinguish between isoforms) were grouped together, and one accession number was arbitrarily considered as representative of each protein group. NSAF7 v. 0.0.1 (https://github.com/tzw-wen/kite/tree/master/windowsapp/NSAF7x64), was used to create the final reports on all detected peptides and nonredundant proteins identified across the different runs and to calculate quantitative label-free distributed normalized spectral abundance factor (dNSAF) values for all detected protein/ protein groups [73]. Raw data and search results files have been deposited to the Proteome Xchange (accession: <u>PXD035300</u>) via the MassIVE repository and may be accessed at ftp://MSV000089877@massive.ucsd.edu with password Kehu-2022-07-13. Original mass spectrometry data underlying this manuscript can be accessed after publication from the Stowers Original Data Repository at http://www.stowers.org/research/publications/libpb-1730.

### Cloning of plasmids

Genomic DNA (gDNA) fragments were amplified using gDNA templates prepared from RHΔ*hx* or RHΔ*ku80*Δ*hx* parasites [74, 75]; a kind gift from Dr. Vern Carruthers, (University of Michigan) using the Wizard Genomic DNA Purification Kit (Cat# A1120, Promega, Madison, WI). Coding sequences (CDS) were amplified using *T. gondii* complementary DNA (cDNA). See Table S2 for primers used in PCR amplification.

#### ptubA1-mEmeraldFP-Pcr1, Pcr2, or Pcr3

The coding sequences for *Pcr1*(TGGT1_274160), *Pcr2* (TGGT1_257370), *Pcr3* (TGGT1_231840) were amplified by PCR using primers S1 and AS1 (*Pcr1*), S2 and AS2 (*Pcr2*), S3 and AS3 (*Pcr3*), digested with *BamH*I and *Afl*II (*Pcr1*) or *Bgl*II and *Afl*II (*Pcr2* and *Pcr3*), and ligated between the *Bgl*II and *Afl*II sites of ptub-mEmerald-TLAP2_231-410aa-T7, which has the same structure as ptubA1-mEmeraldFP-TUBA1[69].

#### ptubA1-mEmeraldFP-Pcr2_V2

The coding sequence for *Pcr2* was amplified by PCR using primers S4 and AS4, and inserted between the *Bgl*II and *Afl*II sites of ptubA1-mEmeraldFP-TUBA1[69] using the NEBuilder HiFi Assembly kit (New England Biolabs, #E2621S).

#### pTKO2_II-mEmeraldFP-Pcr2 (for generation of mEmeraldFP-Pcr2 knock-in parasites)

∼1.9 kb fragments upstream (5’UTR) or downstream (3’UTR) of the Pcr2 genomic locus were amplified from the parasite genomic DNA by PCR using primers S5 and AS5 (5’UTR) and S6 and AS6 (for 3’ UTR) and inserted between the *Not*I-*Xho*I (5’UTR) or *Hind*III-*Nhe*I (3’UTR) sites of the pTKO2-II-mCherryFP [61] using the NEBuilder HiFi Assembly kit. The vector backbone of the resulting vector was used to generate pTKO2_II-mEmeraldFP-Pcr2, into which the CDS for mEmeraldFP-Pcr2, PCR amplified using primers S7 and AS7 from ptubA1-mEmeraldFP-Pcr2_V2, was HiFi assembled between the *AsiS*I and *Rsr*II sites. A linker sequence coding for SGLRS was added in between the Pcr2 and Emerald coding sequences, and the Kozak sequence from the endogenous Pcr2 locus (TATTACCAGTGAAatg) was added to the 5’ end of the mEmerald coding sequence. The backbone of the pTKO2_II-mEmeraldFP-Pcr2 plasmid contains a cassette driving expression of cytoplasmic mCherryFP, to help identify and exclude non-homologous or single homologous recombinants.

#### pPcr2-mNeongreenFP-LIC-DHFR (for 3’ endogenous tagging of Pcr2 with mNeonGreenFP)

was generated by HiFi assembly using three components, including the vector backbone, Exon 6 of *Pcr2*, and a CDS for mNeonGreenFP. The vector backbone was generated from pYFP-LIC-DHFR (a kind gift from Dr. Vern Caruthers, University of Michigan) digested with *Pac*I and *BssH*II. Exon 6 of *Pcr2* without the stop codon was amplified from *T. gondii* genomic DNA with primers S8 and AS8. The CDS for mNeonGreenFP was amplified using primers S9 and AS9 from plasmid pmNeonGreenFP-N1 (a kind gift from Dr. Richard Day, Indiana University) [76].

### Generation of knock-in, endogenously tagged, knockout, complemented, and transgenic parasites

#### 3’ endogenously tagged Pcr2-mNeonGreenFP parasites

RHΔ*ku80*Δ*hx* parasites (∼1 × 10^7^) were electroporated with 40 µg of the endogenous tagging plasmid Pcr2-LIC-mNeongreenFP linearized by *BstZ17*I within exon 6. Parasites were selected with 1 µM pyrimethamine, then cloned by limiting dilution. Five clones were tested with diagnostic genomic PCRs to confirm that the *pcr2* locus had been fused with the CDS for mNeonGreenFP. One clone was further confirmed by Southern blot (Figure 4A)

#### mEmeraldFP-Pcr2 knock-in parasites

RH*ΔhxΔku80* parasites were electroporated with 40 µg of pTKO2_II-mEmeraldFP-Pcr2 linearized with *Not*I and selected with 25 µg/mL mycophenolic acid and 50 µg/mL xanthine. Clones were screened by fluorescence for mEmeraldFP concentration at the apical end of the parasites and for lack of the marker for non-homologous insertion (cytoplasmic mCherry fluorescence), and confirmed with diagnostic genomic PCRs. Two clones (KI-a and KI-b) were further verified by Southern blot and were used in the generation of *Δpcr2* parasites (Figure 4A).

#### Δpcr2 parasites

*mEmeraldFP*-*Pcr2* knock-in parasites were electroporated with 30 µg of pmin-Cre-eGFP_Gra-mCherry [59], selected with 80 µg/mL of 6-thioxanthine, and screened for the loss of mEmerald fluorescence. Clones were confirmed by diagnostic genomic PCRs. Two clones (KO-a and KO-b, derived from KI-a and KI-b, respectively) were further verified by Southern blot (Figure 4A).

#### Δpcr2:mEmeraldFP-Pcr2 complemented parasites

*Δpcr2* parasites were electroporated with 40 µg of pTKO2_II-mEmeraldFP-Pcr2 linearized with *Not*I, and selected with 25 µg/mL mycophenolic acid and 50 µg/mL xanthine.

### Southern blotting

Southern blotting was performed as previously described [59, 61] with the following modifications. To probe and detect changes in the *pcr2* genomic locus in the parental (RHΔ*ku80*Δ*hx*), 3’ endogenously tagged *Pcr2-mNeonGreenFP, mEmeraldFP-Pcr2* knock-in parasites, *Δpcr2* parasites, and complemented parasites, 5 µg of gDNA from each line was digested for hybridization with a CDS probe and a 3’UTR probe. To generate the CDS probe, a CDS fragment was released from the plasmid pTKO2_II-mEmeraldFP-Pcr2 by *Rsr*II digestion, gel-purified, and used as a template in probe synthesis by nick translation. To generate the 3’UTR probe, a fragment specific for the region downstream from the last *pcr2* exon was released from the plasmid pTKO2_II-mEmeraldFP-Pcr2 by *Afl*II and *Hpa*I digestion, gel-purified, and used as a template in probe synthesis. For hybridization with the CDS probe, parasite genomic DNA was digested with *Rsr*II. The predicted *Rsr*II fragment size is 5270 bp for the parental (*i*.*e*., wild-type *pcr2* locus), 2946 bp for the knock-in, and ∼ 9.1kb for the *Pcr2-mNeonGreen* 3’ tagged line. As expected, no signal was detected in the lane with *Δpcr2* genomic DNA when hybridized with the CDS probe. For hybridization with the 3’UTR probe, parasite genomic DNA was digested with *Nco*I. The predicted *Nco*I fragment size is 8197 bp for the parental, 8665 bp for the knock-in, 2151 bp for the knockout, and ∼ 10.9 kb for the *Pcr2-mNeonGreen* 3’ tagged line.

### Generation of rat TgCEN2 and TgGAP45 antibodies

Purified recombinant TgCEN2 (TogoA.00877.b.A2.PW30868) and TgGAP45 (TogoA.17128.a.A1.PW28089) proteins were kindly provided by the Seattle Structural Genomics Center for Infectious Disease (Seattle, WA), and used to inject rats for antibody production (Cocalico Biologicals, Inc). Sera of the immunized animals were harvested for performing the immunofluorescence labeling of TgCEN2 and TgGAP45.

### Plaque assay

One hundred freshly harvested parasites per well were used to infect confluent HFF monolayers grown in 6-well plates. After incubation at 37°C for 9 days, the cultures were rinsed with PBS, fixed with 70% ethanol for 15 min, stained with 2% (w/v) crystal violet, rinsed with PBS, air-dried and scanned using an Epson Perfection V500 photo scanner.

### Wide-field deconvolution microscopy

Unless otherwise noted, image stacks were acquired at 37°C using a DeltaVision imaging station (Applied Precision) with a COOLSNAPHQ2 camera (Photometrics®), and an Olympus 100X oil (NA 1.40 or 1.35), Olympus 60X oil (NA1.40), or Olympus 60X silicone oil immersion lens (NA 1.3). Deconvolution of the image stacks was carried out using Softworx (Applied Precision). Images were contrast adjusted to optimize their display.

### Immunofluorescence assay

For intracellular parasites, *T. gondii*-infected HFF monolayers growing in MatTek glass-bottom dishes (MatTek Corporation, CAT# P35G-1.5-14-C or CAT# P35G-1.5-21-C) were fixed in 3.7% (w/v) formaldehyde in PBS for 15 min, permeabilized with 0.5% (v/v) Triton X-100 (TX-100) in PBS for 15 min, blocked in 1% (w/v) bovine serum albumin (BSA) in PBS for 30-60 min, followed by antibody labeling (see below). Dishes were incubated in primary antibodies for 30-60 min followed by washing and incubation in secondary antibodies for 30-60 min unless otherwise noted. Primary antibodies and dilutions used were as follows: mouse anti-IMC1, 1:1,000 (a kind gift from Dr. Gary Ward, University of Vermont); mouse 6D10 anti-TgMIC2 antibody [77] (a kind gift from Dr. Vern Carruthers, University of Michigan), 1:1,000; mouse anti-TgRON4 (a kind gift from Dr. Maryse Lebrun, Université de Montpellier, France), 1:1,000; rat anti-TgGAP45 (this study), 1:500; rat anti-AKMT, 1:1,000 [7]. Secondary antibodies and dilutions used were: goat anti-mouse IgG Alexa350, 1:1,000 (Molecular Probes); goat anti-rat IgG Cy3, 1:1,000 (Jackson ImmunoResearch, 112-165-167). All immunofluorescence labeling steps were performed at room temperature.

For labeling of TgCEN2 in extracellular *Pcr2-mNeonGreenFP* parasites, parasites were allowed to attach to poly-lysine coated MatTek glass-bottom dishes, permeabilized with 0.5% TX-100 in PBS for 15 min, fixed in 3.7% (w/v) formaldehyde in PBS for 15 min, then blocked with 1% (w/v) BSA for 30min. The samples were then incubated with rat anti-TgCEN2, 1:500 (this study) for 60 min, followed by goat anti-rat IgG Cy3 secondary antibody at 1:1,000 for 60 min.

### Replication assay

Intracellular replication assay was performed as described in [7]. The replication rate was calculated using the method described in [78]. Four independent experiments were performed. In each replicate, the number of parasites in ∼ 100 vacuoles was counted for each strain at each time point.

### Invasion-related assays

To quantify differences in invasion efficiency at the population level, immunofluorescence-based invasion assays were performed and analyzed as previously described [22, 62] with some modifications. ∼5 × 10^6^ freshly egressed parasites were used to infect a MatTek dish of a nearly confluent monolayer of HFF cells. After 10 min incubation on ice and then 1 hr incubation at 37°C, the dishes were washed with PBS and fixed with 3.7% (w/v) formaldehyde for 15 min. To label the extracellular parasites, the samples were blocked with 1% (w/v) BSA in PBS for 30 min and incubated with a rabbit antibody that recognizes the *Toxoplasma* <u>s</u>urface <u>a</u>nti<u>g</u>en 1 (TgSAG1) (a kind gift from Dr. Lloyd Kasper, Dartmouth College) at 1:2,000 dilution for 30 min, followed by goat anti-rabbit IgG Alexa 568 at 1:1,000 dilution for 30 min. To label both extracellular and intracellular parasites, cells were then permeabilized with 0.5% (v/v) TX-100 in PBS for 30 min, blocked with 1% (w/v) BSA in PBS for 30 min, incubated with the same rabbit anti-TgSAG1 antibody at 1:2,000 dilution for 30 min, followed by goat anti-rabbit IgG Alexa 488 (Molecular Probes) at 1:1,000 dilution for 30 min. The samples were imaged using an Olympus 20X (NA 0.70) objective. Ten full field images per sample were collected for each of three independent experiments. Fields were randomly selected using the Alexa 488 channel. Parasites that were labeled with both Alexa 568 and Alexa 488 were scored as uninvaded (*i*.*e*. extracellular), and parasites that were labeled with Alexa 488 only were scored as invaded (*i*.*e*. intracellular).

To analyze the behavior of live parasites during invasion, extracellular parasites were used to infect a MatTek dish of a nearly confluent monolayer of HFF or BS-C-1 cells. The dishes were gently centrifuged at 1,000 rpm (Eppendorf Centrifuge 5910R) for 4min at 10°C, or incubated on ice for ∼10 min, to facilitate parasite settling on the host cells. The culture was then imaged at 37°C at 1 sec (Figure 9 and Video S8) or 1.5 sec (Video S1) intervals to capture invasion events.

To analyze the assembly of the moving junction in invading parasites, extracellular parasites were used to infect a MatTek dish of a nearly confluent monolayer of BS-C-1 cells. After ∼ 10 min incubation on ice and then 4 -9 min incubation at 37°C, the dishes were washed with PBS and fixed with 4% (w/v) formaldehyde for 15 min. To label the portion of the plasma membrane of the parasite that had not yet been shielded by the host cell plasma membrane, the samples were blocked with 1% (w/v) BSA in PBS for 40 min and incubated with rabbit anti-TgSAG1 at 1:100 dilution for 60 min. Cells were then permeabilized with 0.5% (v/v) TX-100 in PBS for 15 min, blocked with 1% (w/v) BSA in PBS for 60 min, incubated with a mixture of mouse anti-TgRON4 at 1:1,000 dilution and rat anti-TgGAP45 at 1:500 for 60 min, followed by goat anti-rabbit Alexa 488, goat anti-mouse Alexa568, and goat anti-rat Alexa647 (Life Technologies-Molecular Probes) at 1:1,000 dilution for 60 min. Parasites that formed a constriction were first identified by DIC and then recorded with fluorescence and transmitted light (DIC) imaging. Three independent experiments were performed for each line in Figure 10. The number of parasite analyzed for each line in each experiment was 16, 21,15 for RH*ΔhxΔku80* (WT); 15, 15, 15 for *mEmeraldFP-Pcr2* knock-in (KI); 12,18,13 for *Δpcr2*; and 16,16,12 for the complemented (Comp) parasite.

### Egress assays

For induced egress assay, parasites were added to MatTek dishes containing a confluent HFF or BS-C-1 monolayer and grown for ∼35 - 40 hours. Cells were washed once and incubated in 2 ml of L15 media containing 1% heat-inactivated bovine calf serum (“L15 imaging media”), which was then replaced with ∼ 2 ml of 5 µM A23187 in L15 imaging media. Images were collected at 37°C on an Applied Precision Delta Vision imaging station equipped with an environmental chamber. To examine host cell permeabilization during parasite egress, 4′,6-<u>dia</u>midino-2-<u>p</u>henylindole (DAPI) was added to L15 imaging media to a final concentration of 0.5 µg/ml and imaged using a 60X (NA 1.40) or 100X (NA 1.40) oil objective with an exposure time of 50 ms through a BFP filter set. To examine actin kinetics in response to calcium signaling, parasites were transiently transfected with the plasmid pmin-actin-Cb-mE [66] (a kind gift from Dr. Aoife Heaslip, University of Connecticut) and imaged on the Delta Vision imaging station at 37°C before and after A23187 treatment.

### Microneme secretion assay

To examine microneme secretion, freshly egressed parasites were harvested, resuspended in 50 *μ*l L15 imaging media, treated for 10 min at 37 °C with 5 µM A23187 or 50 µM BAPTA-AM, placed on ice for 5 min, and centrifuged for 5 min at 2,000rpm (Eppendorf Centrifuge 5415D) to separate the secreted fraction (supernatant) from the parasites (pellet). The supernatant and pellet fractions were analyzed by Western blot as described in [55] with a mouse 6D10 anti-TgMIC2 antibody [77] at 1:2,000 dilution. GRA8 in the pellet was used as a loading control. The mouse anti-TgGRA8 antibody [79] was a kind gift from Dr. Gary Ward, University of Vermont. To normalize A23187-induced MIC2 secretion, background-subtracted fluorescence of the MIC2 band from the supernatant sample was divided by background-subtracted fluorescence of the GRA8 band in the corresponding pellet sample. The amount of secretion with respect to the wild-type parasite was then calculated.

### 3-D motility assay

The 3-D motility assay for *Toxoplasma*, originally developed in [65], was carried out with the following modifications. Parasites were allowed to infect confluent cultures of HFF cells. Just prior to natural egress, extracellular parasites were removed by rinsing the monolayer with L15 imaging media. The infected monolayer was scraped with a cell scraper before passing the suspension through a 27-G needle to release any remaining intracellular parasites. The released parasites were filtered through a 3 µm Nucleopore filter (Whatman), collected by centrifugation, resuspended in L15 imaging media and kept on ice. 7.5 µl Matrigel (Cat#356237, thawed at 4°C overnight) was mixed with an equal volume of the parasite suspension and injected into a flow chamber assembled by sandwiching a 24×60 mm glass coverslip with a 18×18 mm glass coverslip (VWR) using two strips of 0.09 mm thick double-sided tape spaced ∼ 3 mm apart. The chamber was then sealed with VALAP, a mixture of vaseline, lanolin and paraffin (We found that sealing the chamber significantly reduced sample drift), placed onto the heated stage in an OMX-Flex imaging system (Applied Precision Inc.) and imaged with a heated 60X silicone oil immersion lens (NA 1.3) at 37°C by DIC with the following settings: Image stacks were acquired continuously with 12 ms exposure, 1 µm Z-section spacing, and 41 sections per stack. It takes ∼ 1.6 sec to complete the imaging of each stack. 301 stacks were collected for each imaging experiment.

To facilitate detection and tracking of the parasites’ motility using standard analysis tools, we processed the first 151 frames (∼253 sec) of the 3-D time-lapse DIC images to produce positive valued gray-scale images. We first applied a 3-D median filter (footprint nx = ny = nz = 2 px) to reduce noise. Next, we applied a 2-D Laplacian filter which attenuates out-of-focus portions of the cells followed by a 3-D gradient filter of Sobel type for edge detection. As edge detection did not efficiently detect the cell interior, we then applied a dilation and erosion process using a maximum filter followed by a minimum filter (footprint nz = 3 px, nx = ny =13 px). To avoid edge artifacts introduced by the convolution operations, we cropped 2 pixels from all x and y edges before performing tracking.

For detection and tracking of the parasites in the processed image stacks, we used the TrackMate plugin in Fiji [80, 81]. Parasites were identified with the Difference-of-Gaussian (DoG) detector and tracked with the linear assignment problem (LAP) tracker. Undetected or incorrectly detected spots were manually added or deleted before applying the LAP tracker.

The tracking data from both motile and immotile parasites was exported to comma-separated text files (.csv) and data analysis and plot generation were done using RStudio [82]. In Figure 7C-D, a pause event is defined as an event where the parasite speed stays below the immotile speed threshold for ≥7 consecutive frames (∼11 sec) in a track. The threshold is calculated by averaging the maximum speed of the tracks obtained from the immotile parasites.

### Electron microscopy

Extracellular parasites treated with A23187 as described in [8] were spotted onto carbon-coated copper grids and incubated at 22°C for 30 min in a humid chamber. The grids were then inverted onto 50 µL of 0.5% (v/v) Triton X-100 in H_2_O on a Teflon sheet, incubated for 3 min, blotted to remove most of the liquid, inverted on a drop of H_2_O, blotted again, and then inverted on a drop of 2% (w/v) phosphotungstic acid (pH ∼7) for 3 min. Grids were blotted to remove all of the liquid and allowed to dry. The negatively stained samples were imaged at a magnification of 25,000 or 29,000X on a Technai F20 (FEI) at 120 or 200 keV.

## Acknowledgments

We thank Drs. Dewight Williams and David Lowry of the ASU Ewring/Cowley Electron Microscopy Center for advice and support with electron microscopy, Melissa Molina and Clemente Quintero for tissue culture support. This study was supported by funding from the National Institutes of Health/National Institute of Allergy and Infectious Diseases (R01-AI132463) awarded to K.H.

## Supplemental Videos

**Video S1:** Time-lapse microscopy of two *Pcr2-mNeonGreen* 3’ tag parasites invading the host cell. Upper panels: fluorescence. Lower panels: DIC. Time interval: 1.5 sec. Video speed: 6 frames/s. Scale bar: 5 µm.

**Video S2:** Time-lapse microscopy of A23187 induced-egress for wild-type (*WT*), *mEmeraldFP-Pcr2* knock-in (*mE-Pcr2 KI*), *Δpcr2*, and complemented parasites. A23187 was added at the beginning of the movies to a final concentration of 5 µM. Time interval: 1 sec. Video speed: 60 frames/s. Scale bar: 5 µm.

**Video S3:** Time-lapse microscopy of A23187 induced-egress for wild-type (*WT*), *mEmeraldFP-Pcr2* knock-in (*mE-Pcr2 KI*), and *Δpcr2* parasites. A cell impermeant DNA binding dye (DAPI) was present in the culture medium. The permeabilization of the host cell by the parasites can be detected by the DAPI entering the host cell nucleus to label the DNA. Time interval: 5 sec. Video speed: 12 frames/s. Scale bar: 5 µm.

**Video S4:** Time-lapse microscopy of A23187-induced egress for *Δpcr2* and *Δakmt* parasites. Time interval: 1 sec. Video speed: 60 frames/s. Scale bar: 5 µm.

**Video S5:** Time-lapse microscopy of four examples of wild-type parasites moving in 50% Matrigel. The formation of constrictions can be observed during the parasite movement. For each time point, the projection of the 3-D stack was generated by Stack focuser (ImageJ/Fiji). Time interval: 1.6 sec. Video speed: 4 frames/s. Scale bar: 5 µm

**Video S6:** Time-lapse microscopy of 3-D motility in 50% Matrigel for wild-type (*WT*), *Δpcr2*, and complemented parasites. For each time point, the projection of the 3-D stack was generated by Stack focuser (ImageJ/Fiji). Time interval: ∼ 1.6 sec. Video speed: 10 frames/s. Scale bar: 10 µm. See Video S7 for a dynamic display of the tracks of the parasite movement.

**Video S7:** Videos that display tracks of the movement of wild-type (*WT*), *Δpcr2*, and complemented parasites included in Video S6. The movies were generated using the “Trail movie” function in Softworx (Applied Precision, Inc.). Time interval: ∼ 1.6 sec. Video speed: 10 frames/s.

**Video S8:** Time-lapse microscopy of invasions or invasion attempts for wild-type (*WT*), *mEmeraldFP-Pcr2* knock-in (*KI*), *Δpcr2* (pcr2KO), and complemented (Comp) parasites. Figure 9 includes images selected from the timelapses in the leftmost column, as well as those from the 2nd and 3rd *Δpcr2* timelapses. Time interval: 1 sec. Video speed: 6 frames/s. Scale bar: 5 µm.

**Supplemental Figure S1**. Pcr2 sequence and Alphafold prediction of Pcr2 structure. Residues predicted to form alpha-helices (pLDDT>70) are highlighted in green.

**Supplemental Figure S2**. DIC and projections of deconvolved wide-field fluorescence images of an invading *Pcr2-mNeonGreen* 3’ tag parasite (green), in which the moving junction (anti-RON4, red), and the parasite cortex (anti-GAP45, cyan) were labeled by immunofluorescence.

**Supplemental Figure S3**. Ectopically expressed Pcr1 and Pcr3 are found at the apex (arrows) of the *Δpcr2* parasites. The expression of mEmeraldFP (green) tagged Pcr1 and Pcr3 was driven by a *T. gondii* tubulin promoter. The parasites were labeled by an anti-IMC1 antibody (red) to highlight the parasite cortex.

**Supplemental Table S1**. List of proteins identified in MudPIT analysis of immunoprecipitation using GFP-Trap and lysate from a *eGFP-CEN2* knock-in parasite line [55].

**Supplemental Table S2**. List of primers used in this study.

**Supplemental Table S3**. Raw data for graphs in Figure 5A, Figure 6D, Figure 7C, and Figure 10D.

