## Supplementary material for "An apical protein, Pcr2, is required for persistent movement by the human parasite *Toxoplasma gondii*": Figure S1-S3

TgPcr2 sequence and predicted structure

1 MWSLLGFSEESPPEESQDATAASPPTPSHQDVPASPSATEGKEVAGVASS
51 ASPETMADAPSDSPPATPTSGGGFWSFWGASTPAEAEPAATTPVSSTAAA
101 VETPDSGISETPGPSGVSSSAAESGGDEKQKKKKKKESGKGGEAVAEKKK
151 KKKESEKGGEMVAEKKKGESEEMSSKLVSGSDVEAFRARLEQIELQKKAK
201 EEAKVKRKEDEKKQKEEKREQRGREREEREKKKQEKEAARIAELKRLEEE
251 LKRTQELAEELEKQNEERKNKRKGDRGESSSRGSSSSPSSGRSRREASEE
301 RRKAREEREKRRREEQAGLEELLEATERELTETARSLQESIDRQREEREE
351 LERRRQAQVKREMEIRQLAEEATRALEAQQQAARAAAAPAPPGASKGQPP
401 APSLVSPADVKARNFMVALLKRDGGPSLPPPELEQRRLAEPARGETSGAK
451 AEDRQAPELSRKGAAAPTAGRVLAETEVTEGDESMLRPFLAKRGDSVGDE
501 YQEAQYTQQHVIFRGPAFRAFHELVNATKEINHLKEQVEAASRDEEEITT
551 LRERLQTLLDKETKNSESHLERACAIRQALNSISIVFRAAQRRLKTLFFT
601 KLNAAAGAAMPRAVGTSAALRAILGGDSKVTRLGVGLLREVVLRHLRQAW
651 NQWTRYVKPRDAPPGRGAGTGIGTLEDEKATAAIMALREVEDEAEALKKY
701 NEEEKRKLAEVASRLERWRRQLVAAHDAVIHLRENETKRCKSGVREKERG
751 DRRRPRTSSTLDSAEDDAQAPDVLTRLNALVHTEMHRLGLRFENPRAGFG
801 GDSTPHLETRNRDPFRRIPRSFQTENIGHLRLLHGDANGSFPEDRLSSFL
851 ESRTSDNGSVRSARGRGVRRRRSLDENGNKALSGTESDTSSVSESISSGR
901 SSSTSSEGSGALSSSEKRRASSRMSGENPSRRNNTVDEGGRRKKNFDKAC
951 EDAEITHDLAPATRGQPSKSGDQASSQVSVVKGTLPASRIVKTRPKPPPP
1001 PGASEPATFGGGQQRKLQPASGTETSQDDLESAPSTRSSCSGKGPKGTGA
1051 PTGSTVIYADAHHTQASAKAKNAVQAKVKMERDRQRTATIVVGAGPPILA
1101 PKSLVNGPKSGPKPKQ

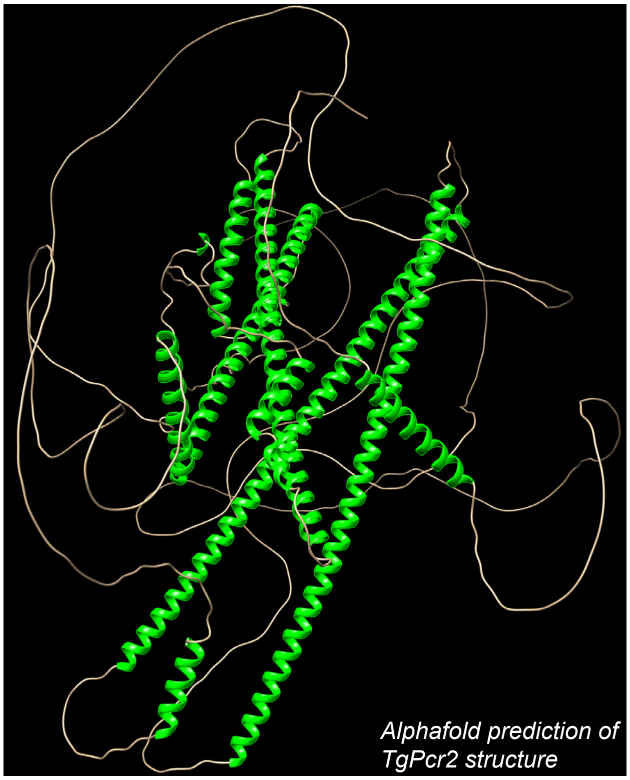


### Figure S2

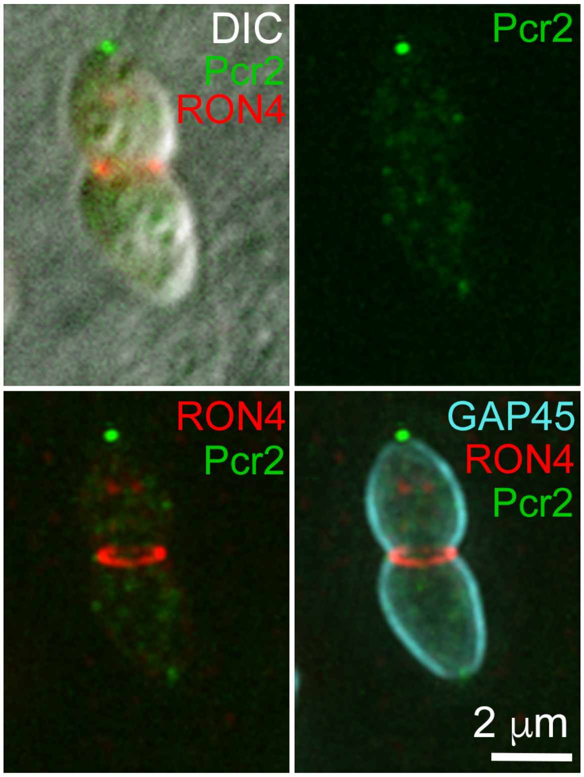


### Figure S3

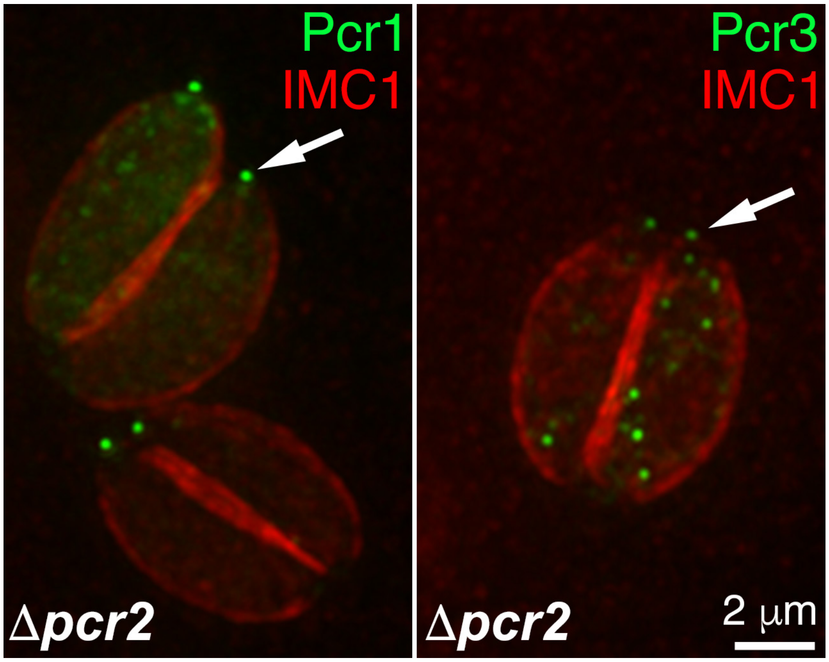
